# Parabrachial *Ntsr1* neurons modulate food intake and anxiety through a projection to the ventromedial hypothalamus

**DOI:** 10.64898/2026.03.03.709382

**Authors:** Jordan L. Pauli, Sekun Park, Rachel R. Felix, Richard D. Palmiter

**Affiliations:** Howard Hughes Medical Institute and Department of Biochemistry, University of Washington, Seattle WA 98195

## Abstract

The parabrachial nucleus (PBN) is an important hub located in the pons that relays sensory signals from peripheral regions. It is genetically diverse and contains many populations that modulate feeding and responses to threatening situations. A small *Ntsr1*-expressing population of neurons was identified that projects selectively to the ventromedial hypothalamus (VMH). The *Ntsr1* neurons are scattered throughout the lateral PBN with a cluster of cells along the border to the nucleus of the lateral lemniscus (NLL) that overlap with *Cck* and *Foxp2* expression. Chemogenetic activation of PBN *Ntsr1* neurons results in *Fos* induction in *Nr5a1* (SF1) and *Bdnf* neurons in the VMH. Activation of PBN *Ntsr1* neurons or their terminals in the VMH reduces food intake after fasting and increases anxiety-like behaviors. In anxiogenic feeding assays, activation of PBN *Ntsr1* neurons increases latency to feed as well as reducing food intake. Photometry showed that PBN *Ntsr1*-neuronal activity increases during anxiogenic situations but is suppressed during food consumption, suggesting a role in threat-induced suppression of feeding. Silencing PBN *Ntsr1* neurons with tetanus toxin light-chain increased food intake and reduced anxiety. These findings reveal a genetically defined PBN to VMH circuit that responds to threats and suppresses feeding behavior.

**Significance:** This study explored a population of *Ntsr1* mRNA-expressing neurons in the parabrachial nucleus (PBN) that project selectively to the ventromedial hypothalamus (VMH). Stimulation of PBN *Ntsr1* neurons and their terminals in the VMH decreased feeding and increased anxiety, results that resemble those achieved by activating VMH neurons, implicating the *Ntsr1* neurons as part of the circuitry that controls feeding and anxiety. Because PBN *Ntsr1* neurons are activated by aversive stimuli, they are posited to help mice suppress feeding in risky environments.

## Introduction

Animals often need to choose between competing internal states, e.g., whether to forage for food and water in a threatening environment. The neural circuits that mediate this balancing act are the subject of numerous studies (1–4). The PBN in the dorsolateral pons is an important hub because it receives sensory input from the spinal, vagal, and trigeminal nuclei along with circulating signals via the area postrema (5–8). Consequently, PBN neurons have been shown to respond to a wide variety of aversive signals and their activation can suppress food intake (9–18). Identification of the post-synaptic targets of PBN neurons that help to mediate behavioral choices in dangerous environments is an ongoing endeavor.

Anatomically, the PBN is bisected by the superior cerebellar peduncle (scp) and is bordered on the dorsal and lateral sides by the nucleus of the lateral lemniscus rostrally and the ventral spinocerebellar tract caudally (19). The neurons in the PBN are genetically heterogenous and can be separated into 2 populations defined by expression of transcription factors, Atoh and Lmx1a/b (20). The projections from the PBN follow central, ventral, and periventricular pathways into the forebrain and a descending tract into the brain stem. The central pathway innervates regions such as the central nucleus of the amygdala and bed nucleus of the stria terminalis, while the ventral pathway innervates regions throughout the hypothalamus; the periventricular pathway innervates regions along the cerebral aqueduct and third ventricle such as periaqueductal gray, periventricular thalamus (PVT), and periventricular hypothalamus, and the descending pathway travels down into the brainstem to innervate areas such as the nucleus of the solitary tract (21–23).

While most PBN neurons send collaterals to multiple brain regions, the *Ntsr1*-expressing neurons (that encode neurotensin receptor 1) are unique in selectively targeting the VMH (23). Like the PBN, the VMH is mainly glutamatergic and genetically heterogenous. It is well known for its roles in sensing and regulating blood glucose (24–27). The VMH has also been shown to regulate food intake and is a well-known target for CCK signaling from the PBN (28–31). Many VMH neurons express *Nr5a1* which encodes steroidogenic factor 1 (SF1); they are known to be important for regulating food intake, anxiety, and hypoglycemia (32–36). Brain-derived neurotrophic factor (BDNF) is also expressed throughout the VMH and affects feeding as well (37–39). The ventral lateral VMH is known for its role in sexual aggression (40–43). While projections from the PBN to the VMH have been noted (44) their functions have received little attention.

SF1 neurons in the VMH become less active during feeding and their artificial activation by chemogenetic means inhibits food intake while promoting anxiety-like behaviors suggesting that these neurons participate in decisions regarding feeding in risky environments (33). The projection of SF1 neurons to the paraventricular thalamus (PVT) has been shown to mediate the inhibition of food intake (34) and BDNF, which is co-expressed in many SF1 neurons, appears to be a major contributor to the decrease in food intake because elevating BDNF expression in the VMH inhibits food intake and knockout of BDNF in the VMH promotes obesity (37). We provide evidence that the excitatory *Ntsr1*-expressing neurons in the PBN project their axons selectively to the VMH where they activate SF1 neurons, are inactive during food consumption, and their chemogenetic activation inhibits food intake and promotes anxiety. Thus, we propose that the parabrachial *Ntsr1* neurons are an important input that drives VMH neuron activity to suppress feeding in risky environments.

## RESULTS

### Anatomical Characterization of *Ntsr1* Neurons in the PBN and Their Projections

To locate the *Ntsr1*-exprssing neurons in the PBN and identify their post-synaptic target regions, *Ntsr1^Cre^*mice were co-injected with AAV-DIO-YFP and AAV-DIO-synaptophysin:mCherry in the PBN to visualize *Ntsr1* cell bodies and axon terminals, respectively (Fig. 1*A*). There is a grouping of Ntsr1 neurons that extend rostrally (Bregma -4.6) and border and partially extend into the nucleus of the lateral lemniscus (NLL) and caudally, around the ventral spinocerebellar tract (sctv) when it appears (Bregma -5.0). *Ntsr1* neurons are also scattered sparsely throughout the PBN with another smaller cluster near the superior lateral and internal lateral regions in the caudal PBN (Bregma -5.25) (Fig. 1*B*). Note that the PBN extends further rostrally and laterally than what is currently shown in commonly referenced atlases as demonstrated by expression of genes such as *Nps, Gal,* and *Ntsr1,* that are classical PBN markers (22, 45).

**Fig. 1.**
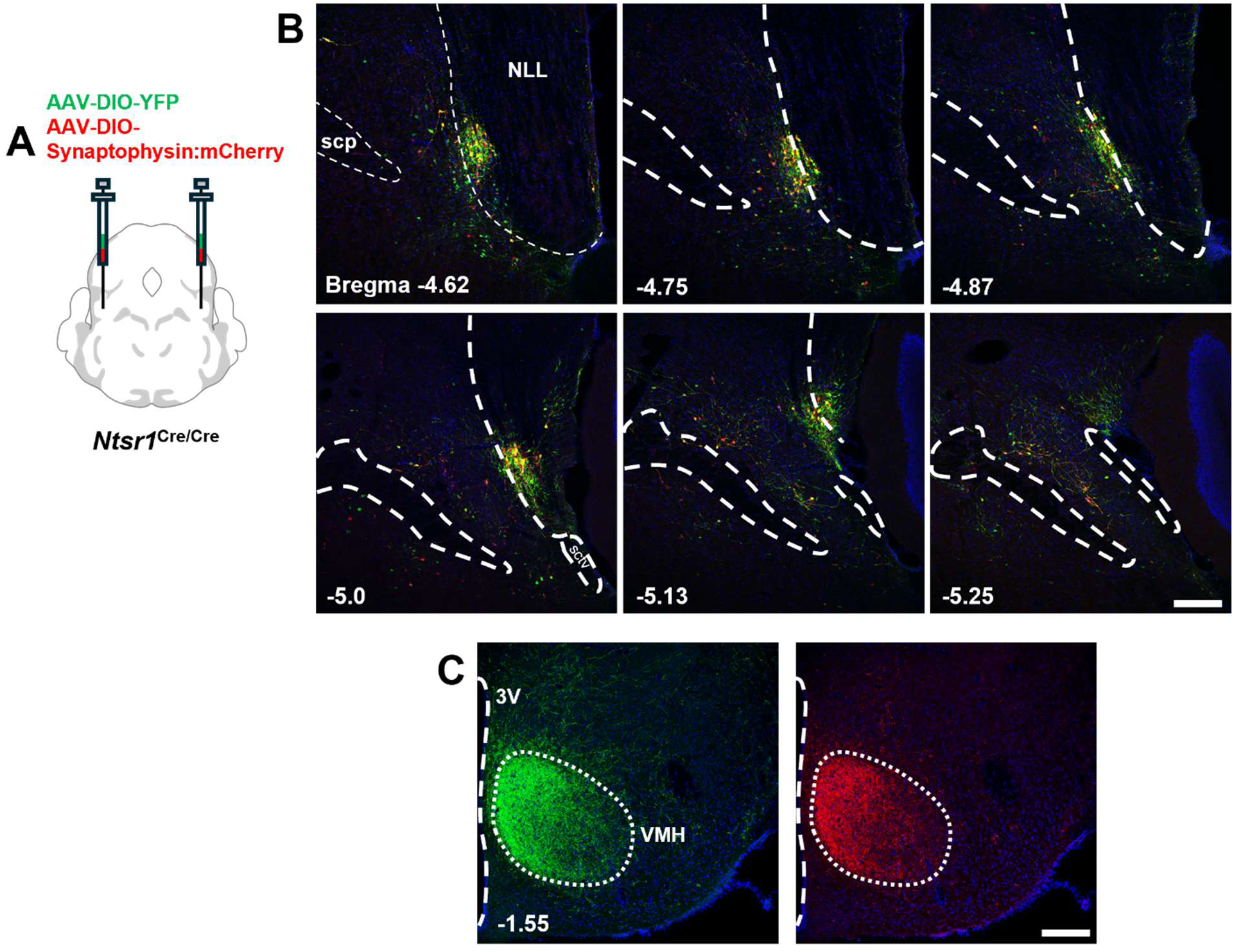
*Ntsr1* neuronal expression pattern in the PBN and their projection to the VMH. (A) Schematic of viral injection of AAV1-Ef1α-DIO-YFP and AAV_DJ_-EF1α-DIO-synaptophysin:mCherry into the PBN of *Ntsr1^Cre/Cre^* mice. (B) *Ntsr1* neurons in the PBN bordering the nucleus of the lateral lemniscus (NLL) in rostral sections and extending throughout the lateral PBN in caudal regions. (C) Innervation pattern of PBN *Ntsr1* fibers (green) and synapses (red) in the VMH. Scale bars: 200 µm. Numbers in lower left indicate bregma level. scp, superior cerebellar peduncle; sctv, ventral spinocerebellar tract; 3V, third ventricle; VMH, ventromedial hypothalamus

The PBN *Ntsr1* neurons were identified previously and were unique among the populations examined because they preferentially innervated the VMH (23). The entirety of the rostral caudal extent of the VMH is innervated from Bregma -1.0 to -2.1 (*SI Appendix*, Fig. S1*A*) with a strong synaptophysin signal with weaker projections to the ventral lateral VMH (Fig. 1*C*). There are also sparse projections (∼20% or lower intensity compared to VMH) to the dorsal medial hypothalamus (DMH), lateral hypothalamus, anterior hypothalamus, and posterior periventricular hypothalamus (*SI Appendix*, Fig. S1*B* and *C*).

### Molecular Characterization of *Ntsr1* Neurons in the PBN

Projections to the VMH from the PBN have been previously identified and were shown to express the neurotransmitter cholecystokinin (29, 32) although the PBN *Cck* neuron projection pathways extend to many other regions besides the VMH (46–48). *Foxp2* has also been identified as a marker for a large portion of the cells in the dorsal regions of the PBN and they also project along the ventral and periventricular pathway to innervate many regions in the hypothalamus (49). To more completely determine the identity of the neurons that express *Ntsr1* in the PBN, fluorescent *in situ* hybridization was used to label *Ntsr1, Foxp2*, and *Cck* (Fig. 2*A*). All three markers are part of the previously identified *Atoh*-derived portion of the PBN (20) and likely have some overlap based on gene expression profiles (23, 50). These markers are more numerous in the rostral regions of the PBN, with the *Ntsr1* population being less than one third the number of the *Cck* and *Foxp2* populations (Fig. 2*B*). *Ntsr1* neurons generally have fewer transcripts per cell (less than 5) than the other markers near the NLL region (Fig. 2*C*). To better visualize the expression of *Ntsr1* and the other markers, a dot map was generated showing the location of *Ntsr1* ‘only’ cells and the location of *Ntsr1* cells that overlap with *Foxp2*, *Cck*, or both in the lateral PBN (Fig. 2*D*). Cells expressing *Ntsr1* mRNA are scattered throughout the entire lateral PBN, with groupings of cells in the rostral lateral and caudal superior regions, replicating the results seen in the tracing experiment. The *Ntsr1* neurons in the rostral PBN that extend into the NLL often coexpress *Cck, Foxp2*, or both, accounting for the dense grouping of cells found there. All three signals are common in the rostral PBN with *Foxp2* and *Cck* more numerous than *Ntsr1* (Full representative PBN expression dot map shown in *SI Appendix*, Fig. S2). Many *Cck* neurons coexpress *Foxp2*, especially in the rostral regions of the PBN but when the whole lateral PBN is considered their overlap falls to around 50%, likely due to the lack of *Foxp2* in the external lateral region. Only 28.9% of *Ntsr1* neurons coexpress *Foxp2* and *Cck* neurons throughout the entire PBN and only 15.1% express all three markers (Fig. 2*E*). Most of the triple labeled cells are localized within the group near the NLL.

**Fig. 2.**
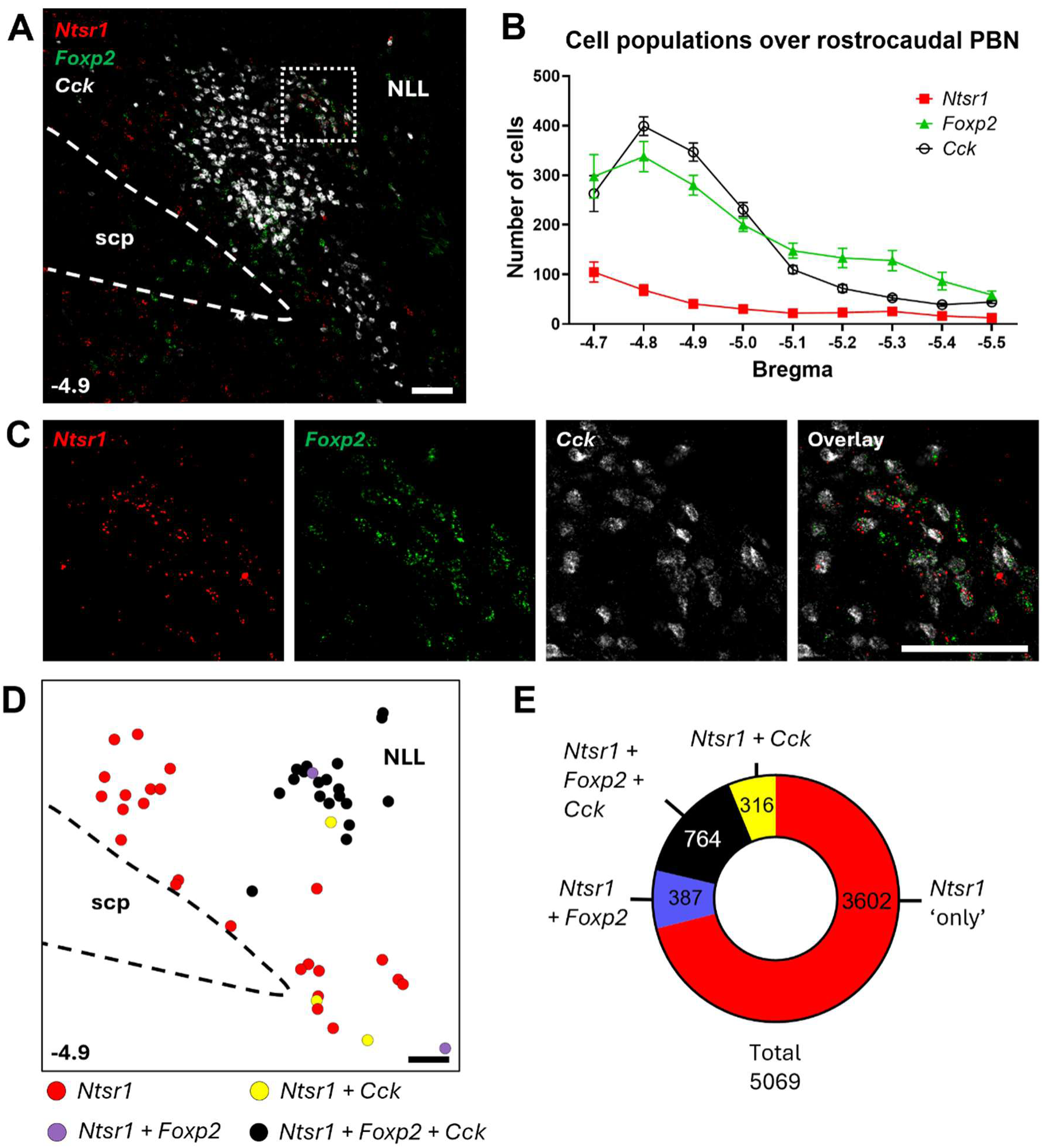
Molecular characterization of PBN *Ntsr1* neurons. (A) Representative image of RNAscope *in situ* hybridization in the lateral PBN for *Ntsr1* (red), *Foxp2* (green), and *Cck* (white). (B) Average number of *Foxp2*, *Cck*, or *Ntsr1* neurons throughout the rostrocaudal extent of the lateral PBN (*n =* 5-6 PBN sections for each group) (C) Magnification of inset in panel A (dotted line) showing individual staining patterns for each probe and their overlap in the PBN neurons that partially extend into the NLL. (D) Representative dot map showing locations of *Ntsr1* neurons along with their overlapping populations in the rostral lateral PBN based on image in panel A. (E) Donut plot showing the overlap between *Ntsr1* (red), *Foxp2* (purple), *Cck* (yellow), and triple-labeled (black) neurons in the lateral PBN. (*n =* 5 whole PBN from 5 animals) Scale bars: 100 µm. Numbers in lower left indicate bregma level. Data are presented as mean ± SEM.

### Chemogenetic Activation of PBN *Ntsr1* Neurons Reduces Food Intake under Both Acute and Chronic Paradigms

PBN neuronal activation has been shown to decrease food intake and promote satiety (9, 12–18). Activating neuronal populations in the VMH that express the transcription factor SF1 and BDNF have also been shown to decrease food intake (33, 37). To determine if the PBN *Ntsr1* neurons modulate food intake, *Ntsr1^Cre/Cre^*mice received AAV injections of Cre-dependent mCherry (as control) or hM3Dq-mCherry in the PBN (Fig. 3*A*) and were activated either acutely or chronically with clozapine N-oxide (CNO). To examine the acute effects, a single dose of CNO was administered (2 mg/kg) intraperitoneally (IP) to fasted mice then a single piece of chow was placed in their cages. Mice with hM3Dq ate significantly less than controls during the 3-hour test (Fig. 3*B*). Several weeks later, mice were moved to special cages for measuring food and water intake constantly (BioDAQ, Research Diets) and CNO (25 µg/ml) was added to the water to examine the effect of chronic CNO administration on eating habits. Over the first 42 hours of chronic CNO exposure, hM3Dq mice ate significantly less food (Fig. 3*C*). CNO exposure decreased the amount consumed during the dark cycle throughout the exposure period, but the greatest effects were seen in the first 2 days (*SI Appendix*, Fig. S3*A*). Mice had no difference at baseline (Fig. 3*D*), but hM3Dq mice ate significantly less food and drank significantly less water during the first 2 days of CNO exposure compared to mCherry controls (Fig. 3 *E* and *F*).The number of feeding bouts also decreased significantly (Fig. 3*G*) along with the amount consumed per bout (Fig. 3*H*), while the time spent feeding per day did not decrease significantly (Fig. 3*I*). The body weight of hM3Dq-CNO treated group also decreased significantly and their food intake was still significantly lower when expressed as a function of their body weight (*SI Appendix*, Fig. S3 *B*-*E*). There were no differences in food consumption between male and female mice (*SI Appendix*, Fig. S4*A*).

**Fig. 3.**
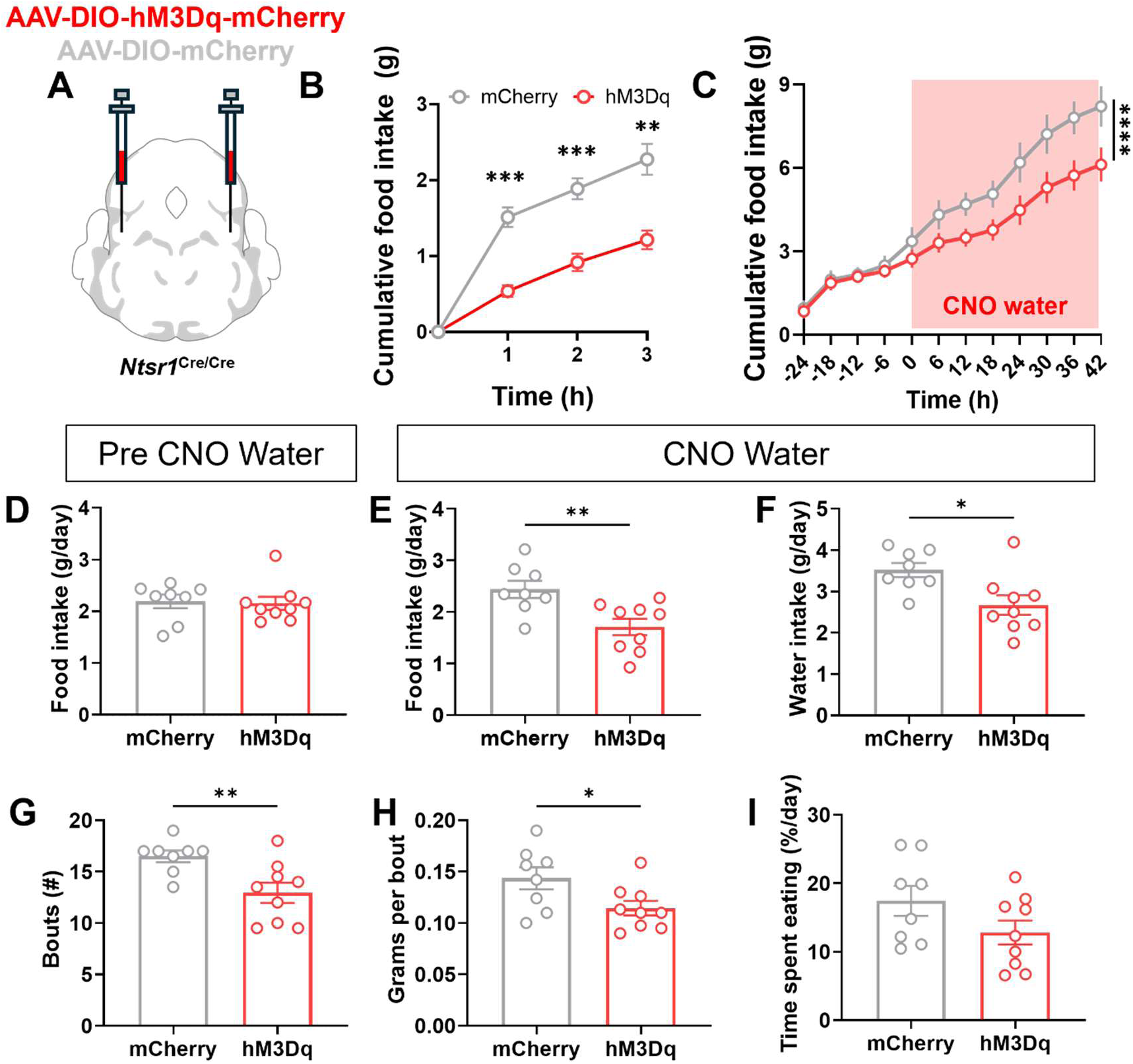
Chemogenetic activation of PBN *Ntsr1* neurons suppresses food and water intake. (A) Schematic of viral injection of AAV1-DIO-mCherry or AAV_DJ_-DIO-hM3Dq-mCherry into the PBN of *Ntsr1^Cre/Cre^* mice (B) Comparison of cumulative food intake after an overnight fast between mCherry and hM3Dq mice. CNO (I.P., 2 mg/kg) was given 30 min before food presentation (*n* = 8 for mCherry, *n* = 14 for hM3Dq, Two-way RM ANOVA, p = 0.0001 for 1 h, p = 0.0002 for 2 h, p = 0.0029 for 3 h) (C) Comparison of cumulative food intake with CNO water (25 µg/ml) between control and hM3Dq mice. Red shade indicates when CNO water was introduced. (Two-way RM ANOVA, p < 0.0001 for mCherry *vs*. hM3). (D) Food intake before CNO water between control and hM3Dq mice (unpaired two-tailed Student’s t test; *p* = 0.8381). (E) Food and F) water after CNO water for the first 2 days (unpaired two-tailed Student’s t test; *p* = 0.0062 for food intake, *p* = 0.0121 for water intake). (G) The number and H) grams per feeding bouts between control and hM3Dq mice during the first 2 days of CNO water (unpaired two-tailed Student’s t test; *p* = 0.0085 for number of bouts, *p* = 0.0362 for grams per bout). (I) Percent of time spent eating each day between control and hM3Dq groups (unpaired two-tailed Student’s t test; *p* = 0.1157). Dots in bar graphs indicate individual animals. *n* = 8 for mCherry, *n* = 9 for hM3Dq in c-i. Data are presented as mean ± SEM.

### Chemogenetic Activation of PBN *Ntsr1* Neurons Reduces Locomotion and Increases Anxiety

The PBN is known to affect locomotion and anxiety (18, 51–53). To explore the effects of activating PBN *Ntsr1* neurons on locomotion and anxiety, open field and elevated plus maze tests were conducted after chemogenetic stimulation. In an open field (Fig. 4*A*), mice with hM3Dq activation showed a significant decrease in distance traveled (Fig. 4*B*) and in the amount of time spent in the center zone compared to controls (Fig. 4*C*) indicating an increase in anxiety. Mice with hM3Dq activation also spent more time in the closed arms and less time in the open arms and center of the elevated plus maze (Fig. 4 *D* and *E*). There was no significant difference in the number of open arm entries or the latency to enter open arms (Fig. 4*F* and *G*). There were no differences between male and female mice in distance or anxiety measures (*SI Appendix*, Fig. S4 *B* and *C*). These results show that the activation of PBN *Ntsr1* neurons induces anxiety and reduces locomotion in novel environments.

**Fig. 4.**
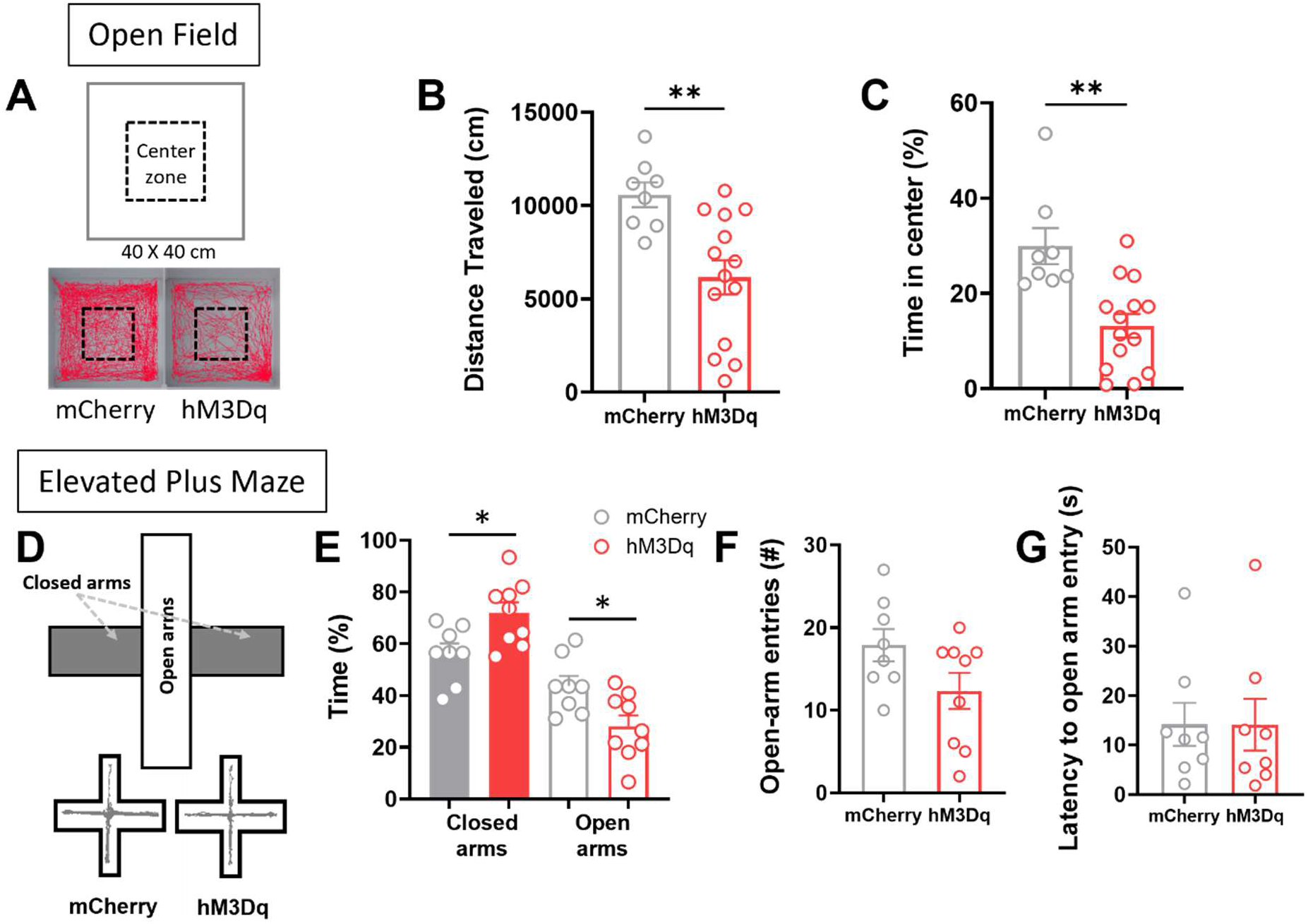
Effect of PBN *Ntsr1* activation on movement and anxiety. (A) Schematic showing the open field and representative traces of mCherry and hM3Dq mice. (B) Comparison of the distance traveled in the open field between mCh and hM3Dq mice (*n* = 8 for mCherry mice and *n* = 14 for hM3Dq mice (unpaired two-tailed Student’s t test; p = 0.0032) (C) Comparison of the time spent in the center of the open field between mCherry and hM3Dq mice (*n* = 8 for mCherry mice and *n* = 14 for hM3Dq mice (unpaired two-tailed Student’s t test; *p* = 0.001) (D) Schematic showing the elevated plus maze and representative mCherry and hM3Dq traces. (E) Time spent in the closed and open arms (*n* = 8 mCh mice and *n* = 9 hM3Dq mice, Two-way repeated measure ANOVA; *p* = 0.0205 for both comparison). (F) Number of open arm entries (*n* = 8 for mCherry mice and *n* = 9 for hM3Dq mice, unpaired two-tailed Student’s t test; *p* = 0.0804). (G) Latency to first open arm entry (*n* = 8 for mCherry mice and *n* = 8 for hM3Dq mice, unpaired two-tailed Student’s t test; *p* = 0.9913). Data are presented as mean ± SEM.

### Chemogenetic Activation of PBN *Ntsr1* Neurons Suppresses Feeding in Anxiogenic Environments

With PBN *Ntsr1* activation impacting both hunger and anxiety, it remained unclear how they might modulate food intake in anxiogenic environments. Activating SF1 neurons in the VMH revealed that mice were less willing to go into novel environments or those associated with shock or predator odor to feed (33). To see if PBN *Ntsr1* neurons also serve a similar function, a novelty-suppressed food-intake test was performed. Overnight fasted mice were placed in a novel open field with a piece of chow fastened in the middle (Fig. 5*A*). Mice with hM3Dq activation took slightly longer to explore the center of the chamber and approach the chow, but the difference was not significant (Fig. 5*B*). They did, however, take significantly longer to take a bite of the food and ate significantly less during the 30-min test (Fig. 5 *C* and *D*).

**Fig. 5.**
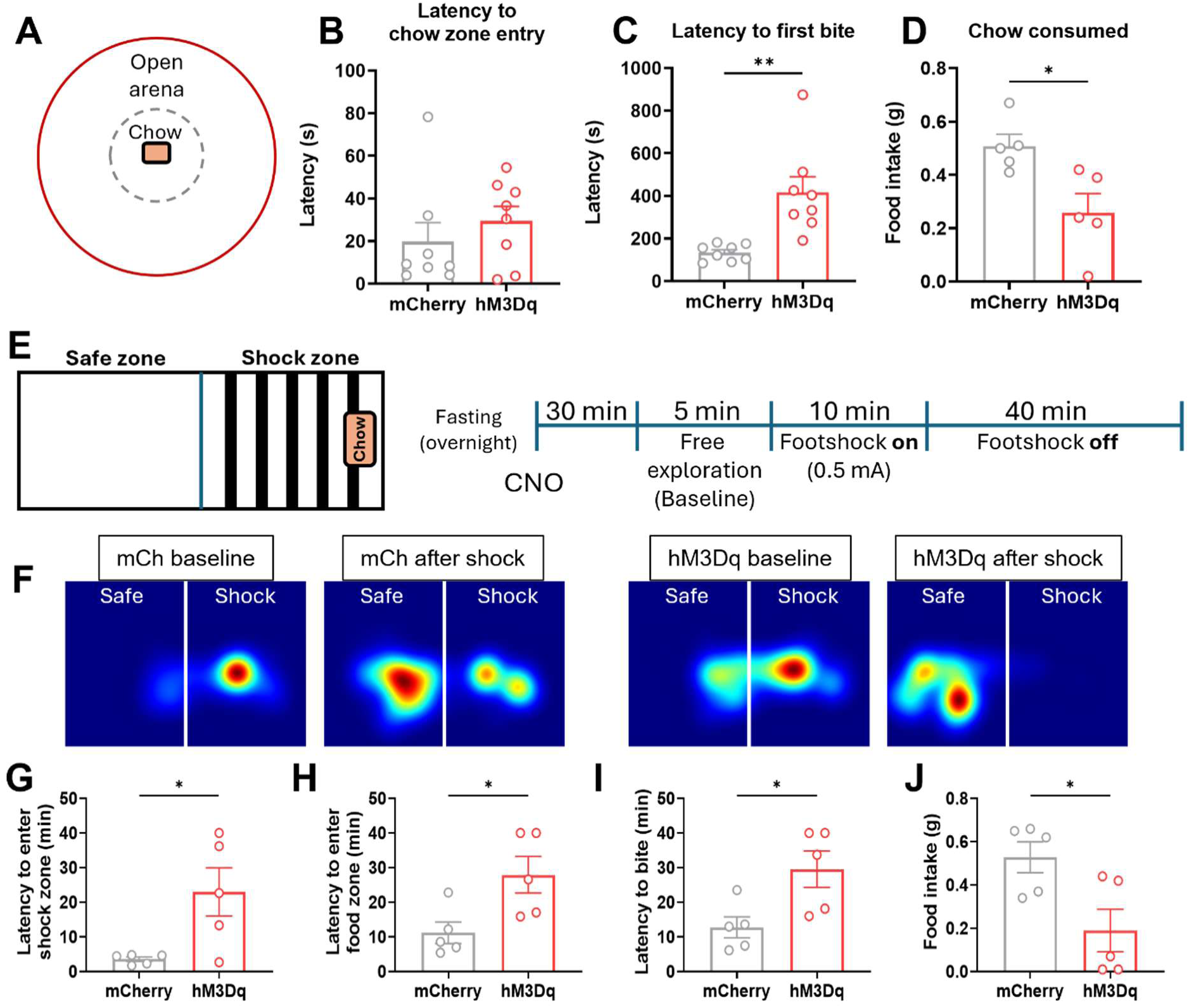
*Ntsr1* activation delays food intake in anxiogenic environments. (A) Schematic showing suppressed feeding test arena. Gray dotted line indicated chow zone. (B-C) Latency to enter the (B) chow zone and (C) first bite in the center of the open field (*n* = 8 for both mCherry and hM3Dq mice, unpaired two-tailed Student’s t-test; *p* = 0.4086 and *p* = 0.0022, respectively). (D) Comparison in amount consumed over the course of the test (*n* = 5 for both mCherry and hM3Dq mice, unpaired two-tailed Student’s t-test; *p* = 0.0178) (E) Left, schematic showing the chamber for the fear-suppressed feeding test. Right, timeline used in the experiment for each mouse is also shown. (F) Representative heatmaps of a single mCherry mouse and a single hM3 mouse showing their preference for each side of the chamber before and after exposure to shock. G-I) Comparison in latency to enter the (F) shock, (G) food zone and (H) first bite after foot shock (*n* = 5 for both mCherry and hM3Dq mice, unpaired two-tailed Student’s t-test; *p* = 0.0241, *p* = 0.0258 and *p* = 0.0242, respectively) (J) Total amount of chow consumed over the entire fear-suppressed feeding test (*n* = 5 for both mCherry and hM3Dq mice, unpaired two-tailed Student’s t-test; *p* = 0.0239). Data are presented as mean ± SEM.

For a fear-suppressed, food-intake test, the mice were placed in a shock chamber with one half of the shock grid covered with a plexiglass floor to create a safe zone and a shock zone (Fig. 5*E*). Chow was fastened to the far region of the shock zone. Overnight-fasted mice were allowed to explore the chamber for 5 minutes, then the shock (0.5 mA) was turned on for 10 minutes. Mice received shocks whenever they stepped onto the exposed grid flooring. After the shock was turned off, mice were allowed to explore for an additional 40 min. Heat maps revealed that mCherry mice returned to the shock side to feed, while hM3Dq mice stayed in the safe zone longer even after the shock was turned off (Fig. 5*F*). Mice with hM3Dq stimulation took significantly longer to return to both the shock zone and the chow zone than controls (Fig. 5 *G* and *H*). They also took significantly longer to take an initial bite of the chow and ate significantly less (Fig. 5 *I* and *J*). These results show that activation of PBN *Ntsr1* neurons suppresses activity and feeding in anxiogenic environments.

The PBN also plays a key role in transmitting painful signals that are caused by injury and noxious temperatures and have been shown to be involved in persistent allodynia (54–59). To determine if the increased avoidance of the painful area in the fear suppressed feeding was a result of a change in sensitivity, heat and mechanical sensitivity were examined. Mice were placed on a hot plate test (*SI Appendix*, Fig. S5*A*) and showed no difference in response latency or number of responses between controls and hM3Dq conditions at 52°C and 55°C (*SI Appendix*, Fig. S5 *B* and *C*). To assess changes in mechanical sensitivity, the von Frey test was used (*SI Appendix*, Fig. S5*D*). There were no changes in mechanical sensitivity between groups even after 4 days of CNO administration (*SI Appendix*, Fig. S5 *E* and *F*). As a result, the PBN *Ntsr1* neurons do not appear to be involved in heat or mechanical sensitivity.

### PBN *Ntsr1* Neuronal Activity Is Suppressed during Feeding

Chemogenetic activation of PBN *Ntsr1* neurons is sufficient to increase anxiety and decrease feeding. Therefore, we hypothesized that the activity of these neurons may decrease during feeding behavior. To examine how PBN *Ntsr1* neurons respond during food intake, GCaMP6m, a Ca^2+^ indicator (60), was expressed in *Ntsr1* neurons and used to monitor their activity by fiber photometry in freely moving mice (Fig. 6 *A* and *B*). Mice were presented with a single piece of chow, and neuronal activity was recorded during approach and consumption phases (Fig. 6*B*). When mice approached the food, Ca^2+^ activity exhibited a sharp increase (Fig. 6 *C* and *D*, and *SI Appendix*, Fig. S6*A*). Mean fluorescence changes (mean ΔF/F) significantly increased during the 2 s following approach compared with the 2 s prior to approach (Fig. 6*E*). Interestingly, these sharp increases in PBN *Ntsr1* neuronal activity during approach were no longer observed after the first consumption bout (Fig. 6*F* and *SI Appendix*, Fig. S6*B*). During consumption, neuronal activity significantly decreased compared with the 2 s preceding consumption onset (Fig. 6 *G*-*I* and *SI Appendix*, Fig. S6*C*). The neuronal activity increased again when the Ca^2+^ activity was aligned to consumption offset (Fig. 6*J* and *SI Appendix*, Fig. S6*D*). Mice were freely moving during recordings, so non-feeding related behaviors were also observed. PBN *Ntsr1* neuronal activity decreased in response to rearing onset but did not show significant changes during grooming behavior (*SI Appendix*, Fig. S6 *E*–*H*).

**Fig. 6.**
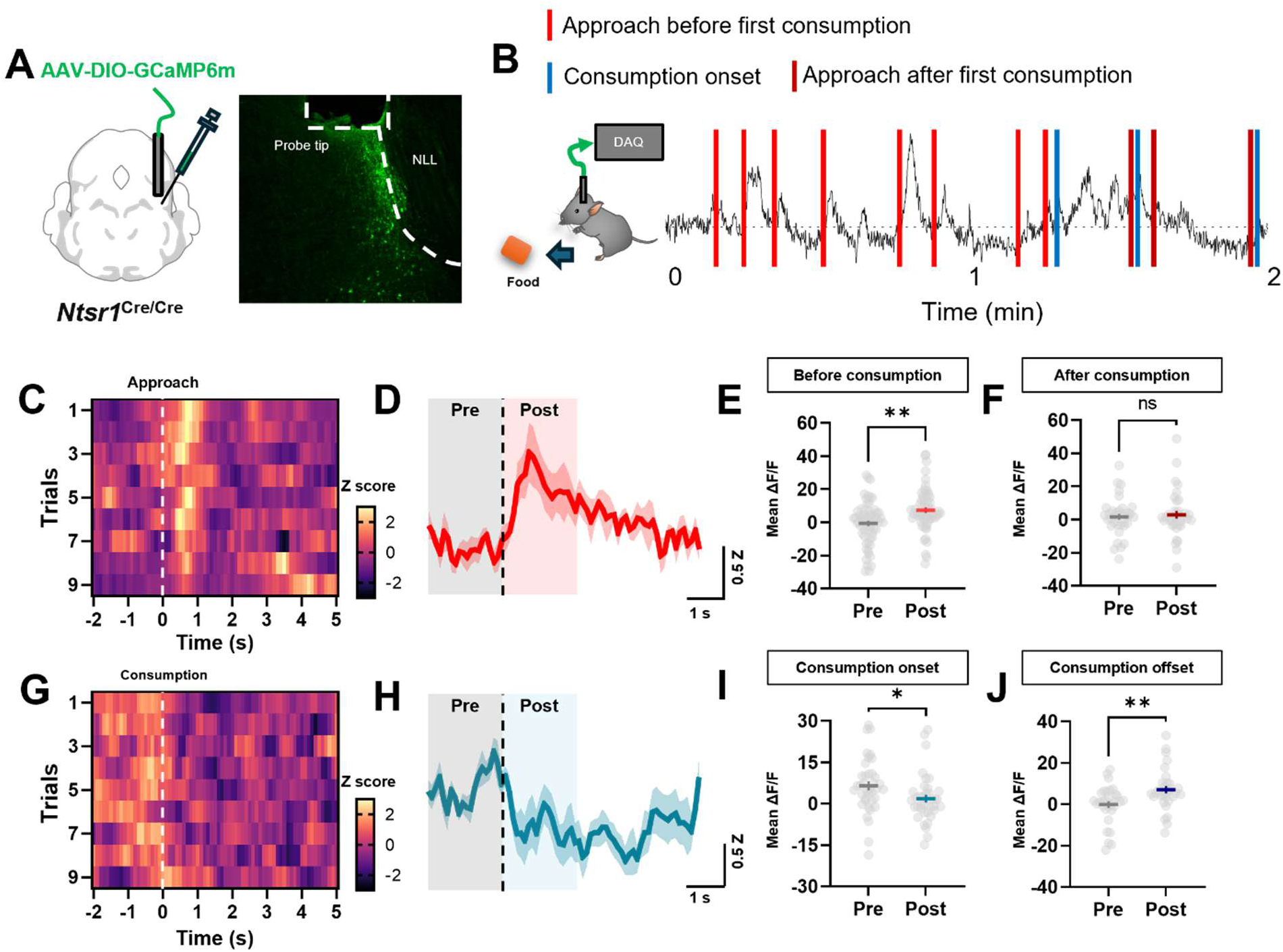
PBN *Ntsr1* neuronal activities in food intake. (A) Schematic of viral injection of AAV-DIO-GCaMP6m and photometry probe unilaterally into the PBN of *Ntsr1^Cre/Cre^* mice along with representative image of GCaMP6m expression and probe placement. (B) Cartoon demonstrating behavioral testing in freely a moving animal with a representative trace of neuronal activity. (C) Heat map of neural activity during approach in a representative animal. (D) Averaged Z score of PBN *Ntsr1* neuronal activity of all mice during approach to the food (*n* = 7). (E) Quantified mean fluorescent changes during approach bouts before first consumption (*n* = 62 bouts in 7 mice, paired two-tailed Student’s t-test; *p* = 0.0028). (F) Quantified mean fluorescent changes during approach bouts after first consumption (*n* = 36 bouts in 5 mice, paired two-tailed Student’s t-test; *p* = 0.7100). (G) Heat map of neural activity during consumption in a representative animal. (H) Averaged Z score of PBN *Ntsr1* neuronal activity of all mice during approach to the food (*n* = 7). (I) Quantified mean fluorescent changes aligned by consumption onset (*n* = 38 bouts in 7 mice, paired two-tailed Student’s t-test; *p* = 0.0280). (J) Quantified mean fluorescent changes aligned by consumption offset (*n* = 31 bouts in 5 mice, paired two-tailed Student’s t-test; *p* = 0.0045). Data are presented as mean ± SEM.

Although chemogenetic activation did not produce detectable changes in pain sensitivity, we next examined whether PBN *Ntsr1* neurons respond to noxious stimuli, similar to other PBN neuronal populations (54–59). To this end, various aversive stimuli were delivered while monitoring Ca^2+^ activity. PBN *Ntsr1* neuronal activity increased significantly in response to tactile noxious stimuli such as tail pinch (*SI Appendix*, Fig. S7 *A*–*C*). Following LiCl administration, which induces visceral pain (14), PBN *Ntsr1* neuronal activity began to increase approximately 5 min after injection and remained elevated throughout the recording period (*SI Appendix*, Fig. S7 *D*–*F*). In addition, noxious somatosensory stimuli such as electric foot shock elicited Ca^2+^ responses in PBN *Ntsr1* neurons, however there was no difference in response to different shock intensities tested (*SI Appendix*, Fig. S7 *G* and *H*).

### PBN *Calca* Neurons activate PBN *Ntsr1* neurons

The *Nts* gene (encodes neurotensin) is broadly expressed in the brain, including the PBN *Calca* neurons that are known to respond to aversive events and suppress feeding (23). Thus, we asked whether activation of *Calca* neurons could potentially regulate *Ntsr1* neurons in the PBN. *Calca^Cre^* mice were treated with either CNO or saline and fresh-frozen brains were collected for fluorescent *in situ* hybridization (*SI Appendix*, Fig. S8*A*). *Calca* neuron activation resulted in a large increase in *Fos* expression throughout the PBN (*SI Appendix*, Fig. S8*B*). The percentage of PBN *Ntsr1* neurons with *Fos* after *Calca* neuron stimulation increased from 6.8% in the control condition to 31.7% (*SI Appendix*, Fig. S8*C*). This result shows that threats that activate *Calca* neurons can activate *Ntsr1* neurons in the PBN, but it does not prove that it is due to intra-PBN signaling or that neurotensin is involved.

### Silencing PBN *Ntsr1* Neurons Increases Food Intake and Reduces Anxiety

To test the necessity of PBN *Ntsr1* neurons in feeding and anxiety, a GFP-fused tetanus toxin light chain (Tetx) was expressed by injecting an AAV with Cre-dependent GFP:Tetx into the PBN (17), while control animals received an AAV with Cre-dependent YFP (*SI Appendix*, Fig. S9 *A* and *B*). Overnight fasting induced robust food consumption during a two-hour period in both groups of mice. The Tetx group showed significantly greater food intake compared with YFP control mice (*SI Appendix*, Fig. S9*C*).

In the open field test, mice expressing GFP:Tetx trended towards more time spent in the center compared with control animals, while overall locomotor activity was not altered (*SI Appendix*, Fig. S9 *D* and *E*). In the elevated plus maze test, however, the anxiolytic effect was significant with Tetx mice spending more time in open arms than control mice (*SI Appendix*, Fig. S9 *F* and *G*). Together, these results suggest that PBN *Ntsr1* neurons normally provide ongoing input to circuits that regulate food intake and anxiety.

### Stimulation of PBN *Ntsr1* Terminals in the VMH Reduces Food Intake and Increases Anxiety

To determine if the projections to the VMH were responsible for the changes in food intake and anxiety, channelrhodopsin (ChR2) was unilaterally expressed in PBN *Ntsr1* neurons, and a fiber was placed ipsilaterally over the VMH to selectively stimulate those terminals (Fig. 7*A*). Previous studies determined that low-frequency stimulation was sufficient to affect behavior in the VMH (33, 61, 62). Activation of PBN *Ntsr1* terminals (5 Hz) in the VMH resulted in a significant decrease in food intake over a 90-min period after an overnight fast (Fig. 7*B*). To assess changes in anxiety, mice were placed in a novel, open field and stimulated and recorded for 15 min. Mice with ChR2 stimulation spent more time around the edge of the arena and less time in the center, indicating an increase in anxiety (Fig. 7*C* and *D*). The latency to enter the center and number of center entries was also affected, but not significantly (Fig. 7*E* and *F*). These results confirm that the VMH is an important target of PBN *Ntsr1* neurons.

**Fig. 7.**
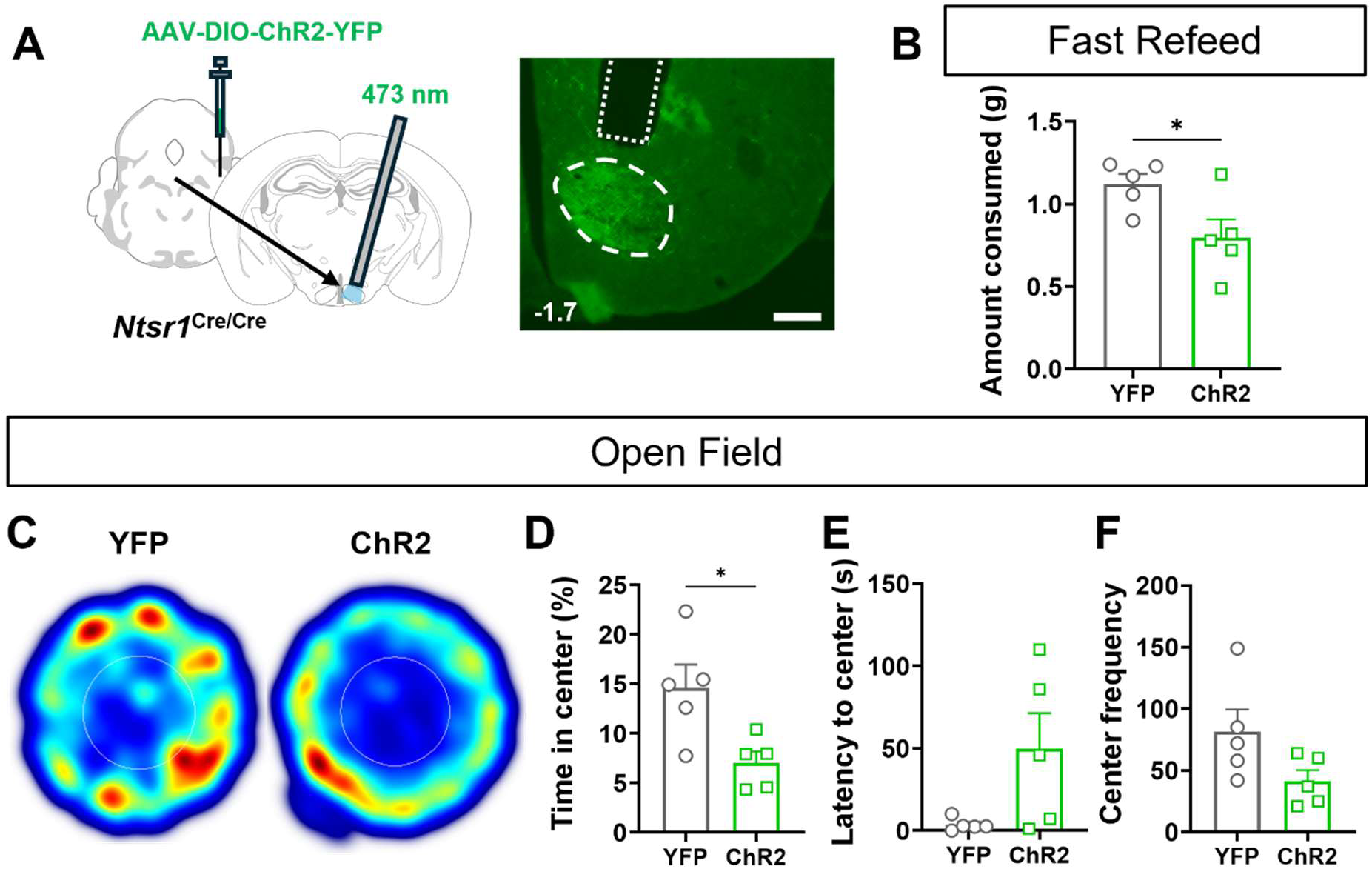
Stimulation of PBN *Ntsr1* terminals in the VMH. (A) Schematic of viral injection of AAV_DJ_-hSyn-DIO-ChR2-YFP into the PBN of *Ntsr1*^Cre/Cre^ mice with fiber placement over VMH and a representative image of terminals and fiber in VMH. (B) Comparison of cumulative food intake after an overnight fast between YFP and ChR2 mice. 5 Hz constant stimulation during the 90 min test (unpaired two-tailed Student’s t test, *p* = 0.0363). (C) Representative heatmaps of a single YFP mouse and a single ChR2 mouse showing their movement patterns over 15 min in a novel open field. (D) Comparison of the time spent in the center of the open field between YFP and ChR2 mice (unpaired two-tailed Student’s t test; *p* = 0.0204). (E) Comparison of latency to first enter the center of the open field between YFP and ChR2 mice (unpaired two-tailed Student’s t test; *p* = 0.0625). (F) Comparison frequency of entries to the center of the open field between YFP and ChR2 mice (unpaired two-tailed Student’s t test; *p* = 0.0870). Scale bar: 200 µm. Number in lower left indicates bregma level. n = 5 for YFP and n = 5 for ChR2. Data are presented as mean ± SEM.

### Chemogenetic Activation of PBN *Ntsr1* Neurons Induces *Fos* in VMH *Nr5a1* Neurons

To determine which VMH neurons are likely post-synaptic targets of the PBN *Ntsr1* innervation, *Fos* expression was assessed alongside *Nr5a1* and *Bdnf* expression, which are abundant in the VMH (33, 35–37). Chemogenetic stimulation was used through AAV delivery of Cre-dependent hM3Dq injected into PBN of *Ntsr1*^Cre^ mice to assess activation of the VMH (Fig. 8*A*). Mice were treated with either CNO or saline and fresh-frozen brains were collected for fluorescent *in situ* hybridization. Chemogenetic PBN *Ntsr1* activation resulted in an increase of *Fos* in the VMH of CNO-treated animals (Fig. 8*B*) resulting in a 106.6% increase in expression (Fig. 8*C*). Both *Nr5a1* and *Bdnf* were expressed throughout the entire VMH with a denser signal for *Nr5a1* in the dorsomedial region and *Bdnf* in the ventral lateral region (Fig. 8*D*). Around 80% of *Nr5a1* and *Bdnf* neurons overlapped with each other across the rostral caudal extent of the VMH (79.4% for *Nr5a1/Bdnf* and 82.62% for *Bdnf/Nr5a1*, Fig. 8*E*). *Bdnf* and *Nr5a1* neuronal populations showed an increase in colocalization with *Fos* after activation with CNO when compared to saline controls (56.94% for *Bdnf* and 52.44% for *Nr5a1*) (Fig. 8 *F* and *G*). A probe for *Cckbr* (CCK_B_ receptor) was also used to look at potential overlap with *Nr5a1* and showed that they also overlapped extensively in the VMH (83.81% for *Nr5a1/Cckbr* and 79.09% for *Cckbr/Nr5a1*, *SI Appendix*, Fig. S10 *A* and *B*). These results indicate that activation of PBN *Ntsr1* neurons is sufficient to activate the *Nr5a1*, *Bdnf*, and, due to the high coexpression with *Nr5a1*, *Cckbr* populations in the VMH. Therefore, PBN *Ntsr1* neurons are likely to activate all of the VMH populations that have been shown to inhibit food intake and promote anxiety (28, 34, 39).

**Fig. 8.**
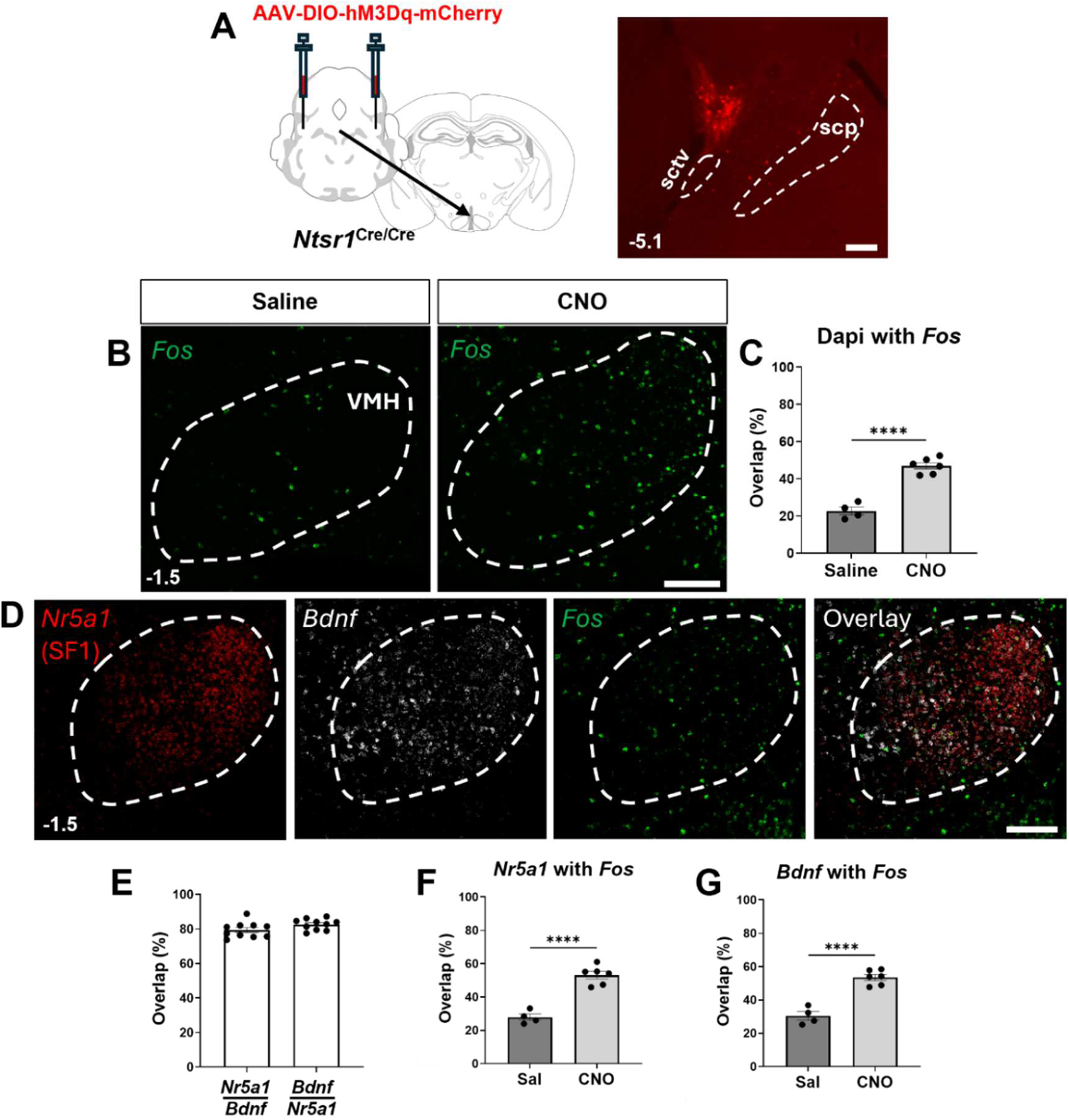
Chemogenetic activation of PBN *Ntsr1* neurons induces *Fos* in VMH. (A) Schematic of viral injection of AAV_DJ_-hSyn-DIO-hM3Dq-mCherry into the PBN of *Ntsr1^Cre/Cre^* mice and a representative image of expression in PBN. (B) Representative images of VMH *Fos* expression in saline vs CNO treated animals. (C) *Fos* expression in the VMH in saline *vs.* CNO treated animals (*n* = 4 for saline, *n* = 6 for CNO, unpaired two-tailed Student’s t-test; *p* < 0.0001) (D) Representative images of *in situ* hybridization in the VMH for *Nr5a1* (red), *Bdnf* (white), *Fos* (green), and their overlap. (E) Percent overlap between Bdnf and Nr5a1 positive populations in the VMH (*n* = 10 for both groups) (F) Percentage of *Nr5a1* cells with *Fos* in the VMH for saline *vs.* CNO treated animals (*n* = 4 for saline, *n* = 6 for CNO, unpaired two-tailed Student’s t-test; *p* < 0.0001) (G) Percentage of *Bdnf* cells with *Fos* in the VMH for saline vs CNO treated animals (*n* = 4 for saline, *n* = 6 for CNO, unpaired two-tailed Student’s t test; *p* < 0.0001) Scale bars, 200 µm; Numbers in lower left indicate bregma level. *n,* number of whole VMH. Data are presented as mean ± SEM.

## Discussion

Evidence is provided that parabrachial *Ntsr1*-expressing neurons provide input to the SF1/BDNF-expressing neurons in the VMH to regulate food intake and anxiety-like behaviors. Activation of *Ntsr1* neurons in the PBN or their terminals in the VMH inhibits food intake by fasted mice in their home environments, novel environments, and in fear-associated environments. Activation of these neurons also promotes anxiety-like behaviors such as avoidance of the middle of an open field and the open arms of an elevated-plus maze. These behaviors resemble the results obtained by activating VMH neurons that express SF1. Photometry experiments revealed that the activity *Ntsr1* neurons decreases as mice consume food, which matches that observed in VMH neurons and reinforces the physiological relevance of the PBN input to the VMH. We confirmed the activation of VMH neurons by showing that chemogenetic activation of Ntsr1 neurons induced *Fos* in *Nr5a1-* and *Bdnf*-expressing neurons in the VMH. Considering that PBN *Ntsr1* neurons project predominantly to the VMH, we suggest that they are an important input to the VMH that redirects attention away from food when mice are in a threatening environment. It has been shown that VMH SF1 neurons project to the PVT to regulate food intake, but that projection does not affect anxiety in an open field (34). We provide evidence that activating *Calca* neurons in PBN, which express CGRP, NTS and several other neuropeptides (23, 50), induces *Fos* in PBN *Ntsr1* neurons. Calca neurons are activated by most threatening conditions as well as cues that predict them (9, 11, 14, 55, 63–65); thus, intra-PBN signaling may complement signaling to other established forebrain targets to inhibit food intake in risky environments. Thus, we suggest that a multi-synaptic PBN^NTS^ ◊ PBN^NTSR1^ ◊ VMH^SF1^ ◊ PVT^VGLUT2^ circuit that participates in redirecting attention away from consumption of food and water when mice are faced with threatening conditions.

The PBN extends further rostrally and laterally than currently depicted in the major atlases. While characterizing various markers in the PBN, *Ntsr1* neurons stood out due to their expression pattern and restricted axonal projections (23). Previous work revealed that *Nps* neurons also lay outside of the current PBN atlas boundaries but are considered part of the PBN due to shared identity, projections, and function with other PBN populations (22, 45). Many PBN *Ntsr1* neurons are near *Nps* neurons in rostral areas and were determined to be a subset of *Cck* neurons (23). In our earlier experiment, *Ntsr1* expression was present in two different glutamatergic clusters, one in the rostral lateral and superior lateral regions, and the other overlapping with *Pax5*, which is expressed in the dorsal lateral regions in the PBN, and likely accounts for the grouping of neurons found in the caudal dorsal regions seen in the *in situ* experiment (See *SI Appendix*, Fig. S2). *Cck* and *Foxp2* were chosen for characterization of PBN *Ntsr1* neurons based on previous analyses of their location and projections (32, 47, 49). Most of the neurons with double or triple labeling are localized in the rostral superior lateral region that borders and extends into the NLL. Other populations that project to the VMH such as *Oprm1* and *Adcyap1* are present in the same region (23). We do not know if the prominent *Ntsr1* cluster in the superior lateral region of the PBN and those scattered caudally have the same projections or functions because our AAV targeting includes both. The precise identity of the *Ntsr1* neurons scattered throughout the PBN outside of that prominent cluster also remains unclear. Although PBN *Ntsr1* neurons have sparse projections to multiple regions such as the LH and DMH (23), the vast majority of the fibers coalesce in the VMH, and it seems likely that the intersection of *Ntsr1* and *Cck* may be useful for specifically targeting this projection.

The VMH plays important roles in glucose metabolism, body weight, food intake, and sexual aggression (32–36, 40–43). The identity of cells in the VMH is also diverse, but studies manipulating SF1 neurons in that region have shown that they are important for regulating food intake and anxiety, with SF1 neurons projecting to the PVT, which was also shown to have a similar function in regards to food intake (33–36). *Bdnf* neurons are also highly expressed in the VMH and were shown to decrease food via input from the arcuate nucleus and projections to the perimesencephalic trigeminal area which affects jaw movement (37). In that and other studies (66), *Bdnf* neurons were shown to have limited overlap with SF1 neurons and represent a distinct subpopulation. However, our *in situ* results show ∼80% overlap between *Nr5a1* and *Bdnf* with most of the *Bdnf* “only” neurons being located in the ventrolateral VMH (VMHvl) compared to the 50% overlap in the dorsomedial VMH (VMHdm) in the previous study. This could be explained by differences in data analysis, staining quality, or that 7-10 sections through the entire VMH were sampled here and were not divided into subregions. PBN *Ntsr1* neuron activation significantly increased activation of *Fos* in *Bdnf* neurons in the VMH. Thus, it is likely that BDNF expression may contribute to the VMH to PVT circuit as well to control food intake and may affect the same circuitry. A third population of neurons expressing the CCK_B_ receptor was of interest because of the partial overlap of *Ntsr1* with *Cck* in the PBN and CCK is a well-known satiety hormone acting through the hypothalamus (30, 67, 68). Silencing *Cckbr* neurons in the VMH has been shown to have a different effects on body weight than silencing of SF1 neurons, suggesting the possibility of different subpopulations in the VMH mediating food intake (69). In the present study, expression of *Cckbr* and *Nr5a1* mRNA also overlapped by ∼80% with *Cckbr* “only” neurons mostly present in the VMHvl, like *Bdnf* expression. Despite potential differences between the results of activating SF1, BDNF, and CCK_B_R neurons, PBN *Ntsr1* neurons likely provide anorexic and anxiogenic inputs to all three types of neurons in the VMH.

The connection between PBN *Cck* neurons and the VMH has been established (28, 31, 32, 47, 29). However, in those cases, the retrograde labeling also included areas near the VMH, or the entire *Cck* population was manipulated to see its effects. *Cck* is expressed across the entire PBN, not just in the rostral superior lateral regions, and projects to many brain regions outside of just the VMH such as the insular cortex, preoptic area, amygdala, and BNST (46–48). Garfield et al (32) showed that Cck neurons in the PBN were important for glucose sensing and exerted their effect via SF1 neurons in the VMH. Blood glucose and other blood hormones were not examined here, but due to the partial overlap between *Ntsr1* and *Cck*, and, the known effects of VMH manipulation, it is likely that the effects of *Ntsr1* PBN stimulation would have similar effects. Another study examined a PBN connection to the VMH that promoted escape-like behaviors (44), but the expression in the PBN was non-specific so the identity of the neurons remains unclear.

Prior studies revealed that activation of SF1 or BNDF neurons in the VMH decreased feeding in mice (33, 37). In our study, PBN *Ntsr1* stimulation also decreased feeding acutely after fasting and chronically through long-term exposure to CNO. In agreement with the downstream effect of VMH neurons, PBN *Ntsr1* neuron stimulation in a long-term feeding assay had a reduced number of bouts of feeding and ate less per bout, along with a lower water intake indicating a reduced motivation to feed even in a safe environment. Anxiety was also shown to be a critical part of SF1 neuron functionality, so mice were examined in basic movement and anxiety tests. PBN *Ntsr1*-neuron stimulation caused mice to spend a decreased time in the center of the open field and the open arms in the elevated plus maze, but they moved less in the open field while distance in the EPM was not different between groups. This decrease in movement is in contrast to increased activity seen in escape-like behaviors induced by non-specific PBN neuron axon terminal stimulation at VMH (44), but that could reflect differences in neuron populations that were stimulated. The decrease in food consumption following an acute fast and the increase in anxiety in an open field were replicated following PBN *Ntsr1* terminal stimulation in the VMH, confirming that the projection there is likely responsible for the behaviors observed when PBN neurons are active.

It has been demonstrated that activation of PBN *Ntsr1* neurons, VMH SF1/BDNF neurons, and glutamatergic PVT neurons promote anorexic signaling. The anxiogenic effect occurs in the PBN *Ntsr1* neurons and the VMH, but the projection to the PVT was not involved (34). It was suggested that the anxiety effect was mild enough in VMH that maybe it was not detected in the open field or that another pathway was responsible. The PVT is known to be involved in anxiety signaling, and that some PBN neurons (but not *Ntsr1* neurons) project directly to the PVT and can promote anxiety (70, 71).

It was determined previously that VMH SF1 neuronal activity is important for the discrimination between the severity of external threats versus the need to feed (33). Mice with PBN *Ntsr1* activation showed less engagement with food in the anxiogenic environments of the novelty and shock-induced fear tests, consistent with the idea that this circuit biases feeding decisions when external threats are elevated. These results are in line with VMH SF1 neuron activation and support a role for PBN *Ntsr1* neurons in integrating aversive and appetite signaling to regulate behavior under threatening or uncertain conditions.

To determine the activity dynamics of PBN *Ntsr1* neurons during food intake, we used fiber photometry to record Ca^2+^ activity of PBN *Ntsr1* neurons in freely moving animals. Consistent with the results of chemogenetic manipulation, we observed a decrease in *Ntsr1* neuronal activity during food consumption. This finding suggests that suppression of PBN *Ntsr1* activity may serve as a permissive change that allows feeding to occur. In contrast, *Ntsr1* neurons were transiently activated when mice initially approached the food; however, this activation was no longer observed after the first consumption bout. This pattern suggests that *Ntsr1* neurons respond to the novelty of the chow and become disengaged once the animals perceive the chow as safe following consumption. These results parallel observations in SF1 and BDNF neurons in the VMH, which show increased activity during stressful situations, whereas feeding behavior is associated with reduced anxiety (32, 33). Further calcium-imaging experiments will help determine whether the activity of PBN *Ntsr1* neurons is also enhanced during more risky situations.

Although silencing PBN *Ntsr1* neurons increased food intake and reduced anxiety in the elevated plus maze, the effect was mild, as silencing showed only a trend toward reduced anxiety in the open field test. This modest effect on feeding and anxiety phenotypes may be explained by previous literature reporting only mild effects on anxiety (34). Given that the *Ntsr1* neurons represent a small neuronal population within the PBN, other PBN neurons regulating food intake and anxiety (12, 18) likely play a more prominent role. Future work using transient inhibition with chemogenetic or optogenetic tools will help define the conditions when *Ntsr1* neuron activity suppresses food intake.

PBN *Ntsr1* neurons were also robustly activated in response to a variety of noxious stimuli, including tail pinch, LiCl induced visceral malaise, and electric foot shock. Although chemogenetic activation of *Ntsr1* neurons did not alter pain sensitivity or hot plate sensitivity, these aversive stimuli strongly activated PBN *Ntsr1* neurons. Given that PBN neurons are activated by nociceptive signals (54–59), and that activation of PBN *Calca* neurons induces *Fos* expression in PBN *Ntsr1* neurons (*SI Appendix*, Fig. S9), *Ntsr1* neurons may be recruited by painful stimuli through this pathway.

## Methods

### Mice

All experiments were approved by the Institutional Animal Care and Use Committee at the University of Washington and were performed under the guidelines of the US NIH Institutes of Health Guide for the Care and Use of Laboratory Animals. *Ntsr1^IRES-Cre^*homozygous mice (referred to as *Ntsr1^Cre^* in the text) were generated as described (23) and backcrossed onto a C57BL/6J background for > 6 generations . Both male and female mice 8 weeks and older were used for experiments. Animals were housed on a 12-hr light cycle at ∼22°C with food and water available *ad libitum* unless otherwise noted.

### Stereotaxic Surgery

Mice were anesthetized with isoflurane and placed on a stereotaxic frame (Neurostar GmbH, Tübingen, Germany). Viruses were injected bilaterally into the PBN (AP -4.9 mm, ML ±1.35 mm, DV 3.4 mm) at a rate of 0.1 μl/min or 0.2 μl/min for 2 min. For tracing experiments, AAV1-Ef1α-DIO-YFP and AAV_DJ_-EF1α-DIO-Synaptophysin:mCherry were injected into PBN. For chemogenetic stimulation experiments, AAV_DJ_-hSyn-DIO-hM3Dq-mCherry or AAV1-hSyn-DIO-mCherry (for control) were injected into PBN, for fiber photometry experiments, AAV1-Ef1α-DIO-GCaMP6m was injected into the PBN and for silencing experiments, AAV_DJ_-hSyn-DIO-GFP:Tetx was injected into the PBN. For optogenetic stimulation, AAV_DJ_-hSyn-DIO-ChR2-YFP was unilaterally injected into the PBN, and a fiber optic cannula was implanted above the ipsilateral VMH (AP -1.35 mm, ML ±1.40 mm, DV 5.1 mm) at a 10° angle. All the viruses were diluted to ∼10^12^ gc/ml scale. Mice were used for experiments >3 weeks after surgery.

### Projection Tracing

For tracing experiments, mice were anesthetized with sodium pentobarbital and phenytoin sodium (0.2 ml, IP) and intracardially perfused with ice-cold PBS followed by 4% PFA. After a 24 h post-fixation in PFA, brains were stored in 30% sucrose for at least 3 days, followed by placement in a mold with OCT, and frozen at -80°C. 35 μm coronal sections were collected for the majority of the brain (Bregma 2.0 to -8.0 mm) on a cryostat (Leica) and stored in sucrose-based cryoprotectant for long term storage at -20°C

For projection staining, sections were washed two times in PBS then incubated in a blocking solution consisting of 3% normal donkey serum and 0.2% Triton-X in PBS for 1 h at room temperature. Sections were then moved to blocking solution containing primary antibodies chicken-anti-GFP (1:10000, Abcam, ab13970), and rabbit-anti-DsRed (1:1000, Takara, 632496) and incubated overnight at 4°C. The following day, sections were rinsed 3 times in PBS and then incubated for 1 h in PBS with secondary antibodies Alexa Fluor 488 donkey anti-chicken (1: 500, Jackson ImmunoResearch, AB 2340375) and Alexa Fluor 594 donkey anti-rabbit (1:500, Jackson ImmunoResearch, AB 2340621). After 3 more PBS rinses, sections were mounted on glass slides and coverslipped with Fluoromount-G with DAPI (Southern Biotech).

Fluorescent images of entire slides were acquired using a Keyence BZ-X710 microscope and higher magnification images were obtained using an Olympus FV-1200 confocal microscope. Images were minimally processed using Fiji to enhance brightness and contrast for optimal representation of the data. All brain section schematics were adapted from Paxinos and Franklin’s Brain Atlas (71). Synaptophysin signal intensity was used to determine synaptic connection. An ROI was placed over brain regions and the average intensity was measured with Fiji after normalization to a non-innervated region to examine innervation strength.

### RNAscope Multiplex Experiments

Mice were anesthetized with sodium pentobarbital and phenytoin sodium (0.2 ml, IP), decapitated, brains were quickly removed, frozen on crushed dry ice, and stored at -80°C. 20 μm coronal sections were cut on a cryostat, directly mounted onto glass slides (SuperFrost Plus), and stored at -80°C. RNAscope fluorescent multiplex assay v2 was performed following the manufacturer’s guidelines. Probes used were: *Ntsr1* (#422411-C1), *Foxp2* (#428791-C3), *Cck* (#402271-C2), *Nr5a1* (#445731-C1), *Bdnf* (#424821-C2), *Fos* (#316921-C1, C3), *Cckbr* (#439121-C2), *Calca* (#578771-C3).

Images were captured on the Keyence at 20x magnification in a 3×3 grid centered on the PBN. Images of the four-channel sets were stitched together using Fiji and subtracted from one another using the image calculator function to remove background auto-fluorescence. Images from all animals were arranged rostral to caudal and aligned with one another based on section anatomy and probe staining patterns. Sections that were too far rostral or caudal were removed before analysis. Images were adjusted and regions of interest were drawn in Fiji and then transferred to the open-source software QuPath for analysis. The subcellular detection function in QuPath was used to define transcript staining, and DAPI signals with 4 or more associated puncta were positive for *Ntsr1* and 8 or more for the other probes. For the PBN, images across the animals were aligned with each other based on section anatomy and probe appearance and sections that were too far rostral or caudal were excluded from analysis.

### Dot Map Generation

Images of the entire right side of the PBN from a representative animal were chosen to make a map. Using the same analysis thresholds as above, QuPath was used to generate an excel file with the X and Y coordinates of every cellular detection. A custom R script was used to read the coordinates and generate a dot to represent the location and identity of each cell. The fiber tract anatomy and location ROIs were copied from section histology.

### Behavior Analysis

For all behavioral studies, multiple cryostat sections through the PBN region were obtained at the end of the experiments to assess the quality of the targeting and level of expression based on fluorescence signals. Mice that did not meet criteria were eliminated from further analysis. For example, in photostimulation experiments, 7 out of 12 mice were excluded based on tissue damage and fiber placement.

### Photostimulation

After recovery from surgery, mice were acclimated to dummy cables attached to the implanted fiber optic cannulas for 3 consecutive days. For food intake and open field studies, blue light stimulation was delivered using a 473-nm laser (5 Hz, 5 ms; LaserGlow) controlled by a waveform generator (Agilent). Light power was adjusted to 10 ± 1 mW at the tip of the fiber optic cannula.

### Food Intake

Mice were given IP saline injections for three days prior to experiment start to habituate them to handling and injections. Individually housed mice had their weight recorded and then were fasted overnight by having their food removed and their cages changed to ensure no leftover food was in the bedding. The following morning, mice were weighed and given CNO (2 mg/kg, IP). Thirty min later a single piece of chow was weighed and placed in the cage. The chow was removed and weighed every hour for 3 hr. For optogenetic stimulation experiments, a piece of chow was weighed and secured to a Petri dish with a metal twist tie. After 20 min of habituation to the empty Petri dish, the chow containing Petri dish was placed in the home cage, and photostimulation was delivered. Food intake (decrease in weight of chow pellet minus crumbs in the dish) was measured at 90 min.

### Open Field

Mice were given an injection of CNO (2 mg/kg, IP) and placed back in their home cages for 30 min. Mice were then placed facing the edge in a novel white plexiglass arena measuring 40×40 cm and allowed to explore for 30 min. For optogenetic stimulation experiments, mice were placed facing the edge of a 40-cm diameter round open field arena and photostimulated for 15 min. Behaviors were recorded using a Logitech camera and scored using Ethovision (Noldus).

### Elevated Plus Maze

Mice were given an injection of CNO (2 mg/kg, IP) and placed back in their home cages for 30 min. Mice were then placed in the center of the elevated arena (dimensions) and allowed to explore for 15 min. Behaviors were recorded using a Logitech camera and scored using Ethovision for locomotion and time spent in open and closed arms.

### Long-term Food Intake

Mice were individually housed in BioDAQ recording chambers (Research Diets) and allowed to habituate to the recording chambers for 3 days. Mice were given CNO water (25 µg/ml) on day 3. The CNO water was replaced with freshly made CNO water every 3 days. Cumulative food and water consumption along with various feeding parameters were collected for another 6 days with the BioDAQ software.

### Hot Plate

Mice were injected with CNO (2 mg/kg, IP) 30 min before being placed on a hot plate (Bioseb) at either 52°C for 1 min or 55°C for 30 sec. Behaviors were recorded with a Logitech camera and latency to hind paw lick or flick, and the number of responses were hand scored afterwards.

### Von Frey

The von Frey assay was performed using the ascending method. Each animal was placed in an 11.5 cm by 7.5 cm chamber with a wire mesh floor. For baseline measurements, mice were injected with saline and allowed to habituate to the chamber for 30 minutes per day for two consecutive days. On experimental days, mice received CNO (2 mg/kg, IP). Von Frey filaments were applied to the plantar surface of each hind paw up to five times. Testing began with a 0.16 g filament and continued until two consecutive filaments elicited a paw withdrawal response in three or more out of 5 applications. If an animal responded or did not respond 3 times before reaching the fifth application, the remaining applications for that filament were discontinued. Paw withdrawal thresholds were determined for each hind paw and then averaged for analysis.

### Novelty Suppressed Feeding

Mice were fasted overnight in their home cages and given a dose of CNO in the morning (2 mg/kg, IP). A single piece of chow was weighed and affixed to the bottom of a circular piece of plastic flooring with a metal twist tie to prevent the animal from moving it to the edge of the chamber. Thirty minutes after injection, mice were placed facing the edge of a novel red circular arena. Mice were allowed to explore the arena and chow for 30 min. A fresh chow pellet was provided for each trial. Behavior was recorded by a webcam (Logitech) and latency to eat was manually scored. The locomotion was analyzed with Ethovision XT15.

### Fear-suppressed Feeding

Mice were fasted in their home cages overnight. In the morning, mice were given a dose of CNO (2 mg/kg, IP). 30 min after injection, mice were placed in a shock chamber that had a clear plexiglass floor covering half of the shock grid to serve as a “safe” zone. A large piece of chow was affixed to the far side of the shock grid with a metal tie to prevent the mouse from moving it during the test. Mice were allowed to explore the chamber for 5 min before the shock grid was turned on (0.5 mA) for 10 min. After the 10-min shock period, the shock remained off for a subsequent 40 min during which the mice could explore the shock zone and consume chow. Mice were recorded by video, and their behavior was scored with Ethovision.

### Fiber Photometry

Mice were connected to the patch cord and habituated for 5 min per day for 3 consecutive days before the recording session. On the recording day, the patch cord was connected between the data acquisition device (RZ10, Tucker-Davis Technologies) and the fiber photometry probe (400 µm diameter, Doric Lens), implanted on the mouse head 10 min before recording. Video recording was synchronized with the fiber photometry signals using the recording software (Synapse, Tucker-Davis Technologies) with a sampling rate of 1 kHz for both the 473 nm excitation and 405 nm isosbestic signals.

Fiber photometry data were extracted and analyzed using a customized MATLAB script based on code provided by Tucker-Davis Technologies. Based on stimulus onset times (e.g., food approach, food consumption, tail pinch), isosbestic-subtracted fluorescence signals were calculated as ΔF/F = (F – F_405_)/F_405_ within a window from −2 s to +5 s around each event and down sampled to 10 Hz. To compare calcium activity across animals, ΔF/F values were converted to Z-scores [Z = (F − F_mean_)/F_std_]. The area under the curve (AUC) was calculated and compared between the 2 s pre- and 2 s post-stimulus periods.

### Data Analysis

All data were analyzed using Prism 10 (GraphPad) software. Data are presented as the mean +/- the standard error of the mean (SEM). The asterisks in the figures represent the p values as follows: p < 0.05*, p < 0.01**, p < 0.001***, p < 0.0001****.

## ACKNOWLEDGMENTS

Special thanks to Susan Phelps, Lucy Aanastas, Kit Mandeville, Raina Yang, Hutch Clarke, and Alexis Rose for maintaining the mice used in these studies and assisting with behavioral experiments. We appreciate the comments from lab members and reviewers.

## Author contributions

J.P., S.P., and R.P. designed research; J.P., S.P., and R.F. performed research; J.P., S.P., and R.F. analyzed data; and J.P., S.P., and R.P. wrote the paper.

---

**Fig. S1.**
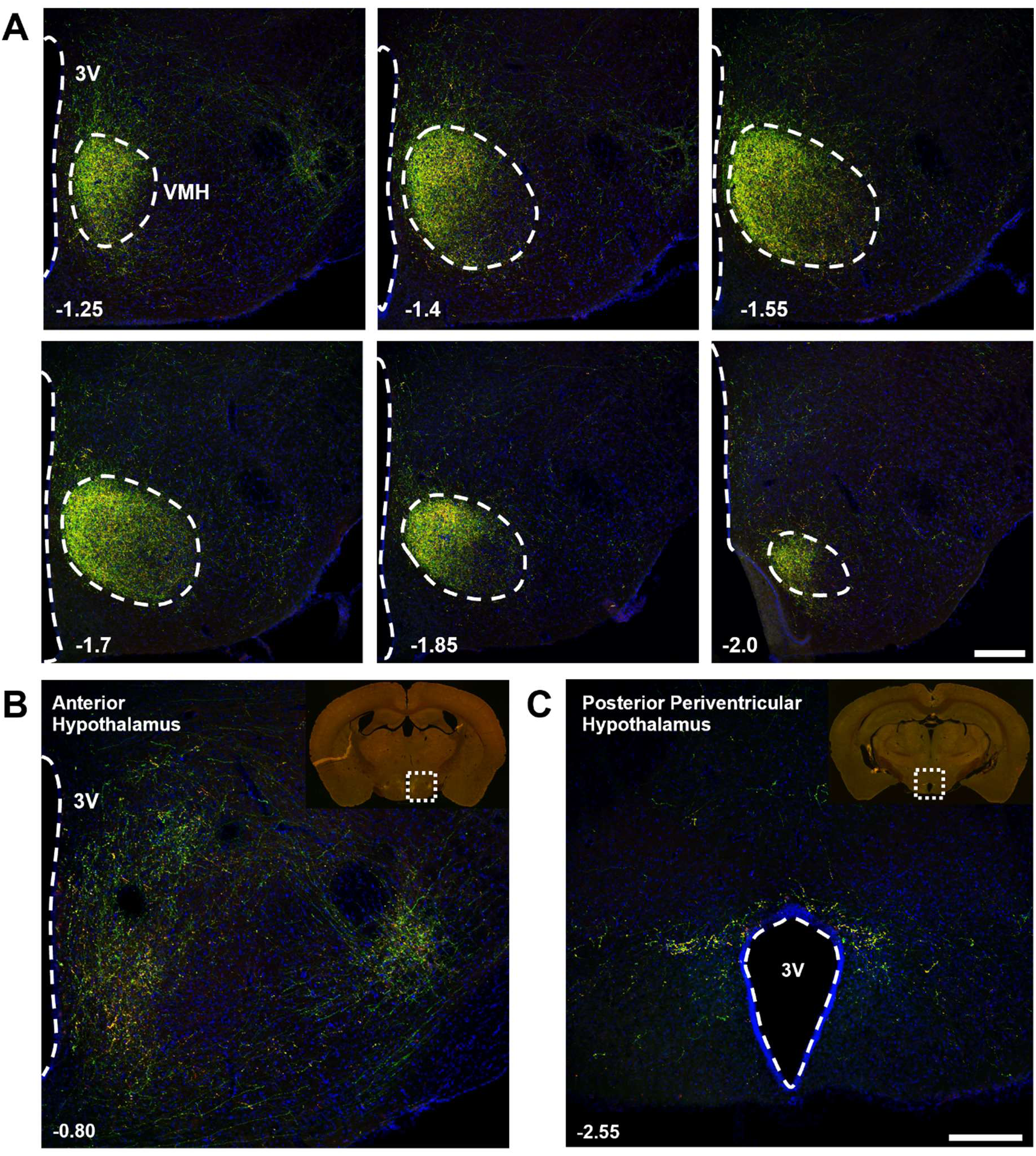
*Ntsr1*^PBN^ neuronal projections in the hypothalamic regions. (A) Images showing the pattern of PBN *Ntsr1* innervation throughout the VMH. (B) Images of PBN *Ntsr1* innervation in the anterior hypothalamus and C the posterior periventricular hypothalamus. (pictures in A-C were taken at the same settings; the average synaptophysin (red) pixel intensity in B and C was 19.84% and 19.80%, respectively, compared to that in **A**. (*n =* 8 hypothalamus sections from 2 mice) Scale bars: 200 µm. 3V, third ventricle; VMH, ventromedial hypothalamus Numbers in lower left indicate bregma level.

**Fig. S2.**
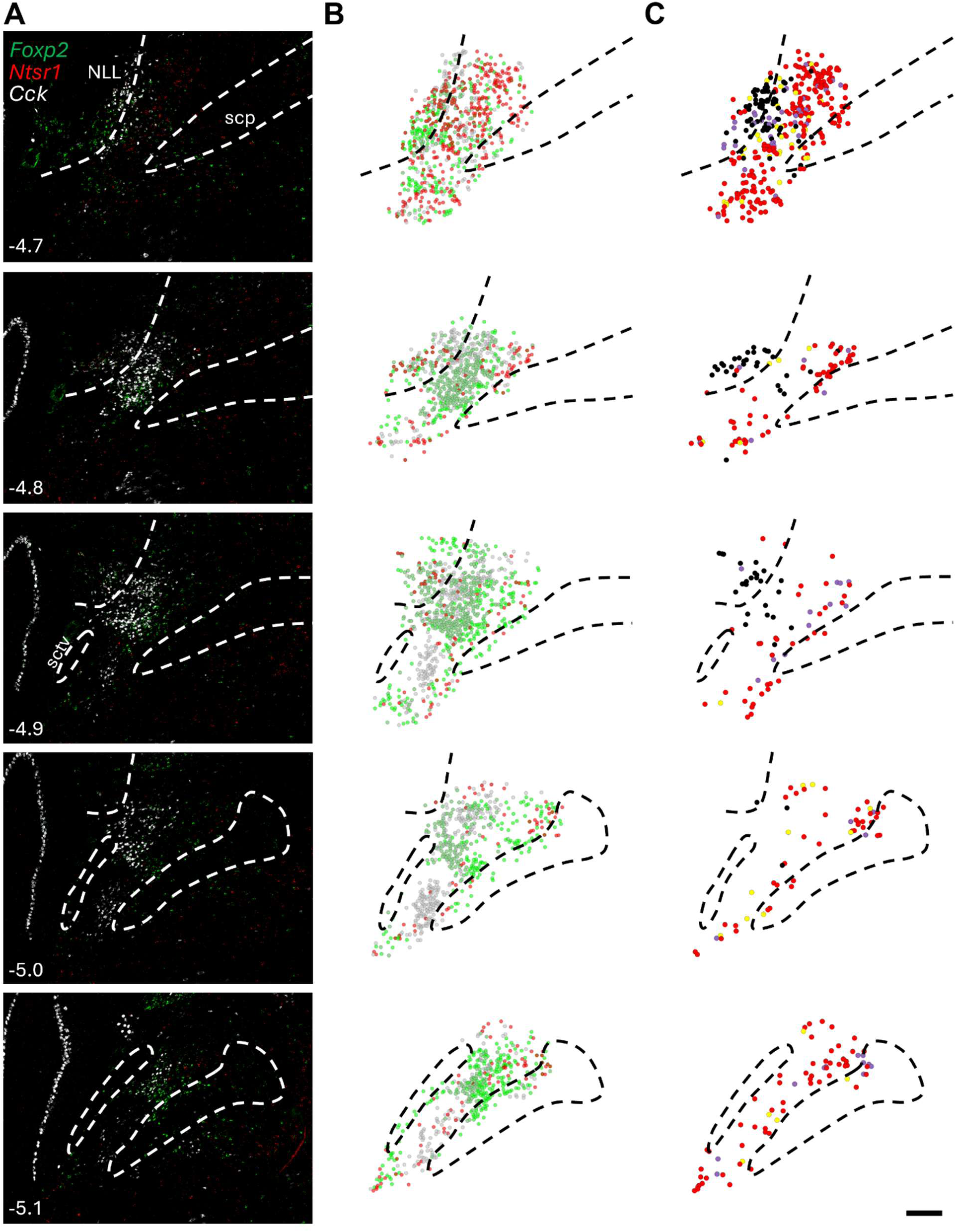
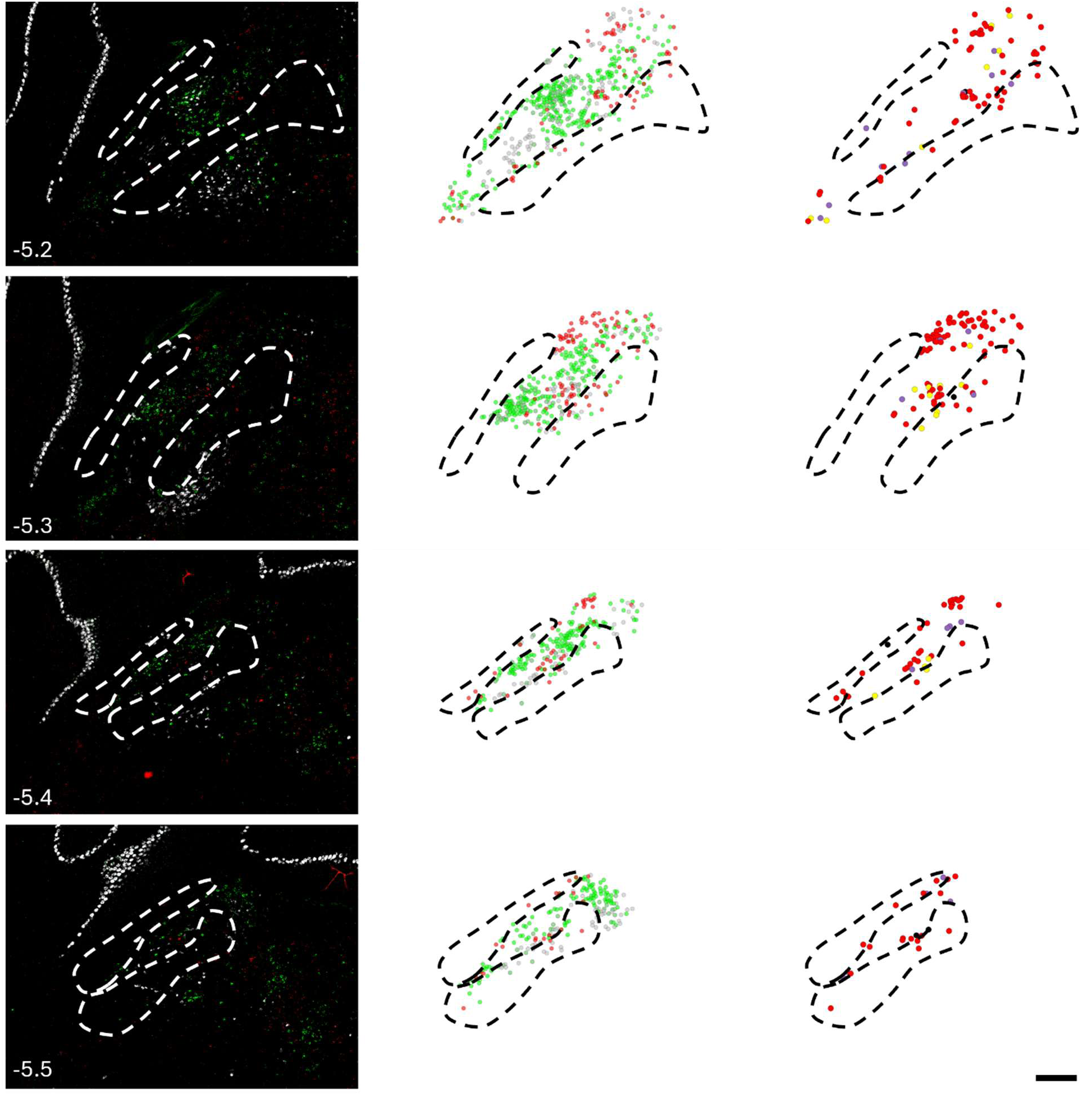
Spatial characterization of PBN *Ntsr1*, *Cck*, and *Foxp2* throughout PBN. (A) Representative images of the left PBN of one animal showing RNAscope *in situ* staining of *Foxp2*, *Ntsr1*, and *Cck*. (B) Dot map of the same PBN sections showing *Foxp2* (green), *Ntsr1* (red), and *Cck* (grey) probes location based on the lateral PBN only. (C) Dot map of PBN *Ntsr1* (red), *Ntsr1* and *Cck* (yellow), *Ntsr1* and *Foxp2* (Purple) and triple labeled (black) cells. Scale bars: 200 µm. Numbers in lower left indicate bregma level.

**Fig. S3.**
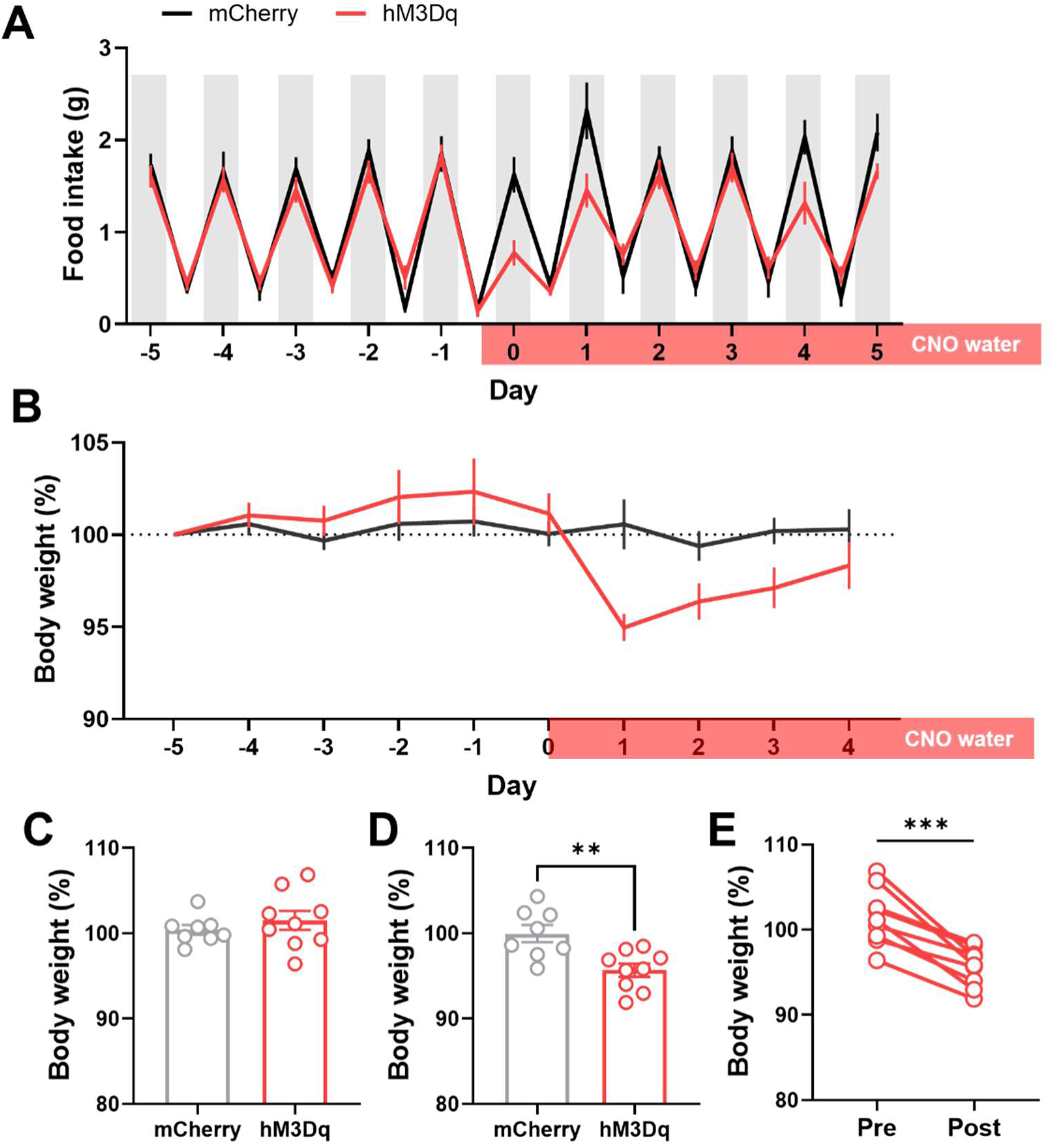
Patterns of food intake over the entire chronic feeding timeline. (A) Graph showing food intake over 5 days before and after CNO water in mCh and hM3Dq mice. Red bar indicates CNO water periods and grey bars show dark cycles (*n* = 8 mice for mCherry and *n* = 8 mice for hM3Dq). (B-E) Comparison of body weight changes based on baseline (day -5) over the chronic CNO exposure feeding experiment before and after introduction of CNO water. Quantification of body weight (C) before CNO water, (D) after CNO water and (E) individual animals in hM3Dq group (*n* = 8 mice for mCherry and *n* = 9 mice for hM3Dq, unpaired two-tailed Student’s t-test, *p* = 0.4092 for C; *p* = 0.0036 for D; paired two-tailed t-test, *p* = 0.0002 for E. Data are presented as mean ± SEM.

**Fig. S4.**
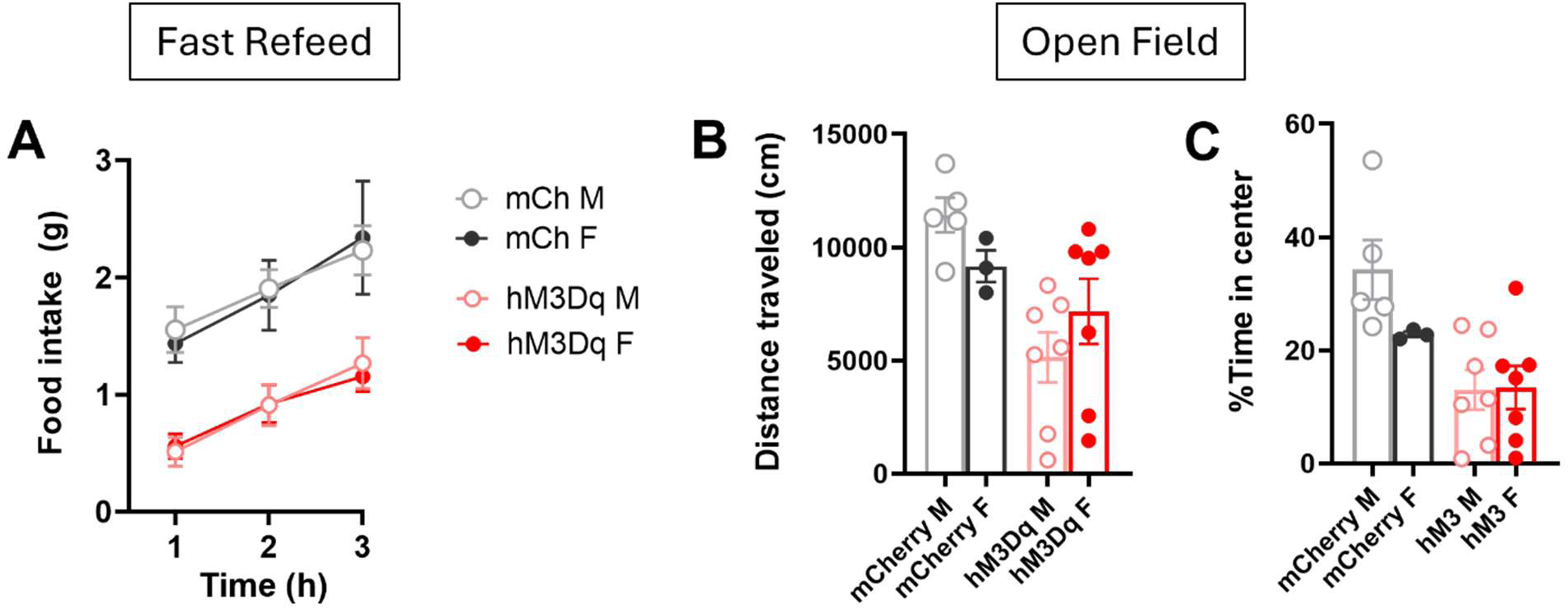
Feeding and movement are the same in both sexes. (A) Comparison in food intake after an overnight fast between female and male mCherry and hM3Dq mice (*n* = 5 and 3 for male and female in mCherry group, Two-way RM ANOVA, *p* = 0.9473; *n* = 7 for both male and female in hM3Dq group, Two-way RM ANOVA, *p* = 0.9299) (B) Comparison in distance traveled in the open field between female and male mCherry and hM3Dq mice (*n* = 5 and 3 for male and female in mCherry group, *p* = 0.0959; *n* = 7 for both male and female in hM3Dq group, unpaired two-tailed Student’s t test; *p* = 0.2865) (C) Comparison in percentage of time spent in the center of the open field between female and male mCherry and hM3Dq mice (*n* = 5 and 3 for male and female in mCherry group, *p* = 0.1550; *n* = 7 for both male and female in hM3Dq group, unpaired two-tailed Student’s t test; *p* = 0.9428) Data are presented as mean ± SEM.

**Fig. S5.**
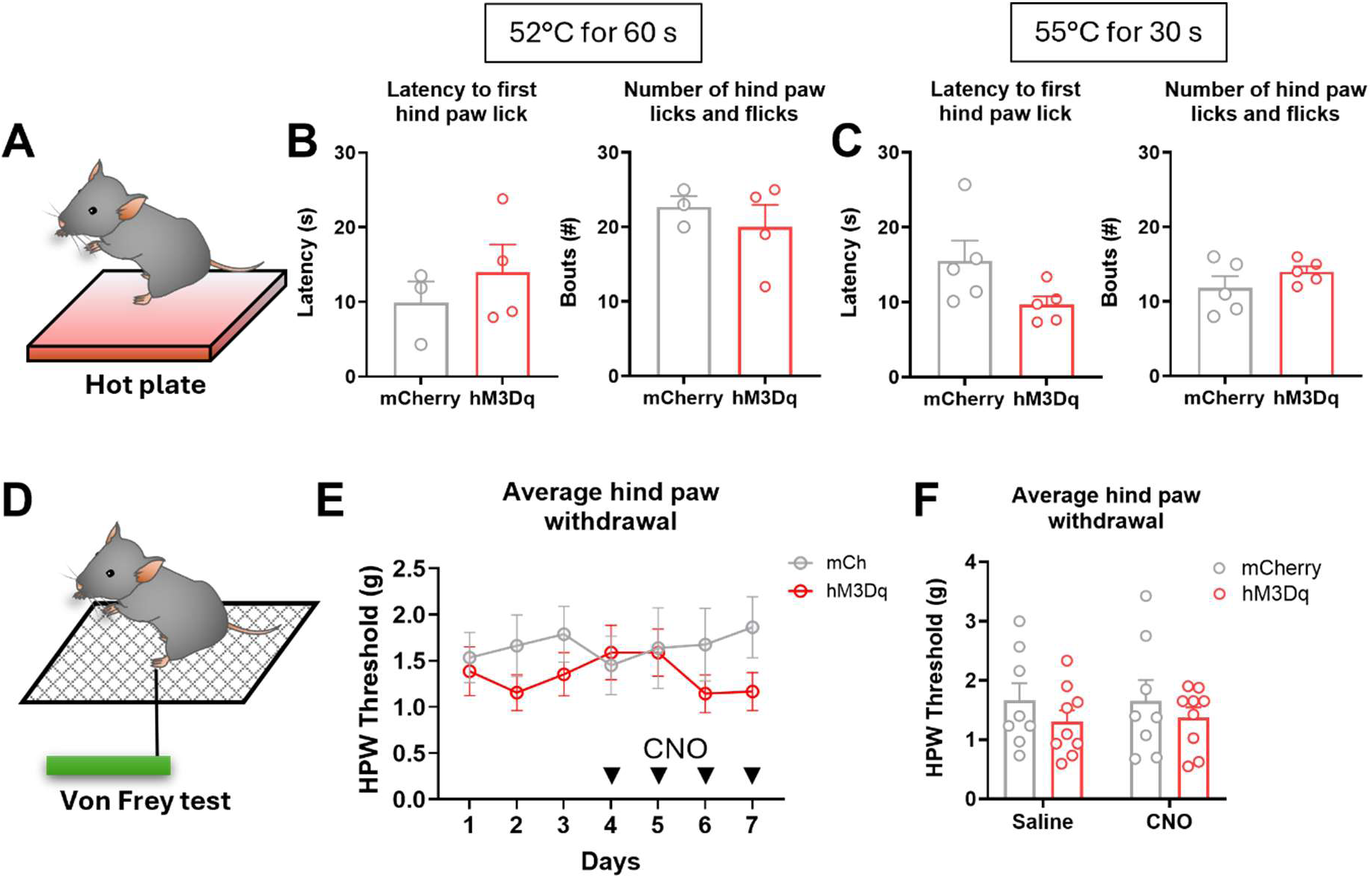
PBN *Ntsr1* neuronal activation has no effect on heat or mechanical sensitivity. (A) Illustration showing hot plate test. (B) Comparisons in latency to first hind paw lick and number of hind paw licks and flicks in a 52 °C hot plate (*n* = 3 for mCherry mice and *n* = 4 for hM3Dq mice, unpaired two-tailed Student’s t-test; *p* = 0.4486 and *p* = 0.5056) (C) Comparisons in latency to first hind paw lick and number of hind paw licks and flicks in a 55°C hot plate (*n* = 5 for mCherry mice and *n* = 5 for hM3Dq mice, unpaired two-tailed Student’s t-test; *p* = 0.0864 and *p* = 0.2426) (D) Illustration showing the schematic showing von Frey filament application to the hindpaw. (E) Paw withdrawal threshold for 7 days. CNO was given on days 4-7 and is marked by arrows (*n* = 8 mCherry for and *n* = 9 for hM3Dq mice Two-way RM ANOVA *p* = 0.3394 for mCherry vs hM3Dq). (F) Bar graphs showing the total averaged hind paw withdrawal thresholds during saline and CNO applications (*n* = 8 for mCherry and *n* = 9 for hM3Dq mice (Two-way RM ANOVA *p* = 0.5340 for Saline, *p* = 0.6782 for CNO). Data are presented as mean ± SEM.

**Fig. S6.**
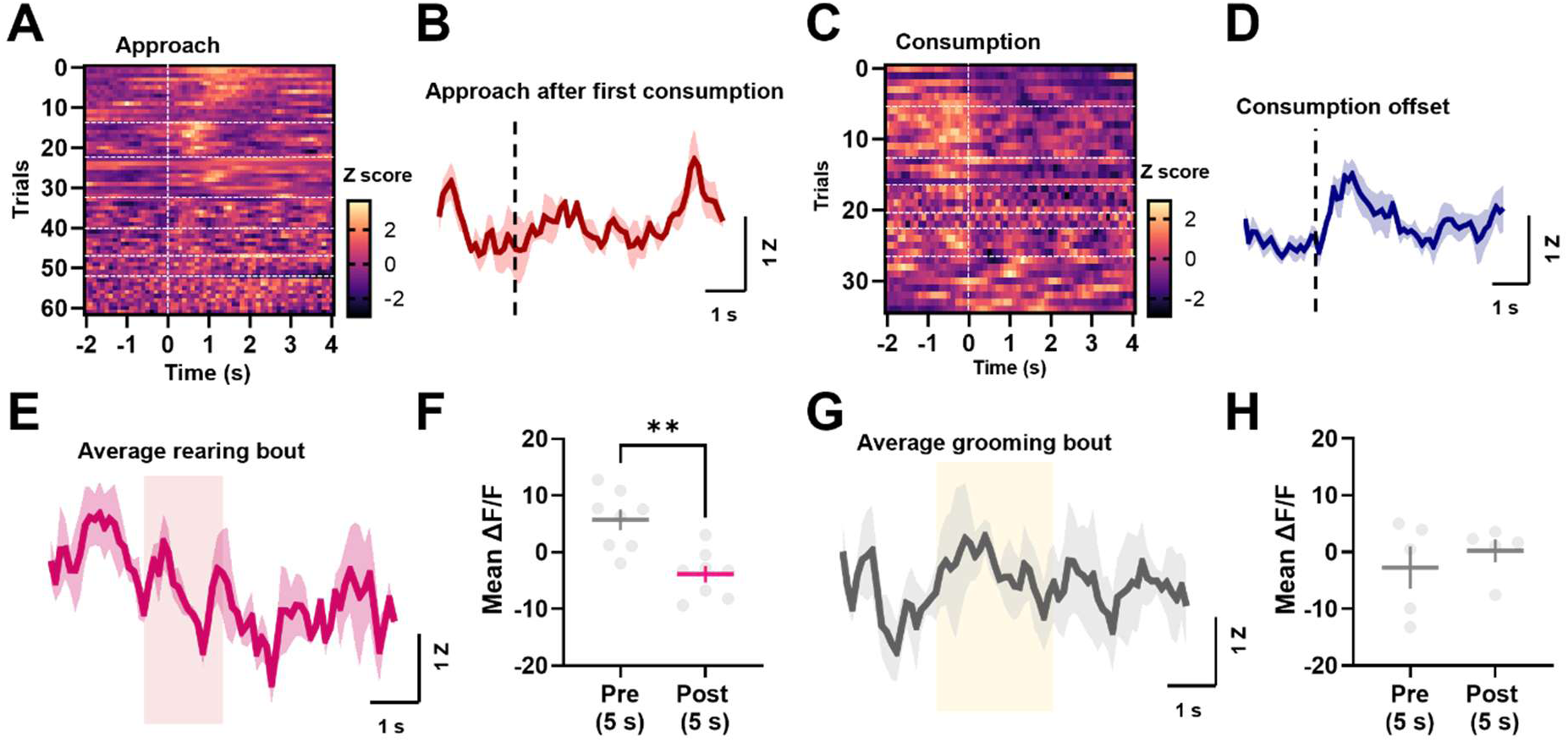
PBN *Ntsr1* neuronal activities during food intake. (A) Heat map of neural activity across approach behaviors from all mice. White dotted horizontal lines indicate individual mice (*n* = 61 trials in 7 mice). (B) Average trace of neural activities during approach behaviors after the first consumption (*n* = 5 mice). (C) Heat map of neural activity across consumption behaviors from all mice. White dotted horizontal lines indicate individual mice (*n* = 34 trials in 7 mice). (D) Average trace of neural activities after consumption behaviors aligned by consumption offset (*n* = 5 mice). (E) Average trace of rearing behaviors during food intake experiment (*n* = 3 mice). (F) Mean fluorescence during the pre 5 s and post 5 s of rearing onset (*n* = 8 bouts in 3 mice, paired two-tailed Student’s t-test; *p* = 0.0040). (G) Average trace of grooming behaviors during food intake experiment (*n* = 3 mice). (H) Mean fluorescence during the pre 5 s and post 5 s of grooming onset (*n* = 5 bouts in 3 mice, paired two-tailed Student’s t-test; *p* = 0.5527). Data are presented as mean ± SEM.

**Fig. S7.**
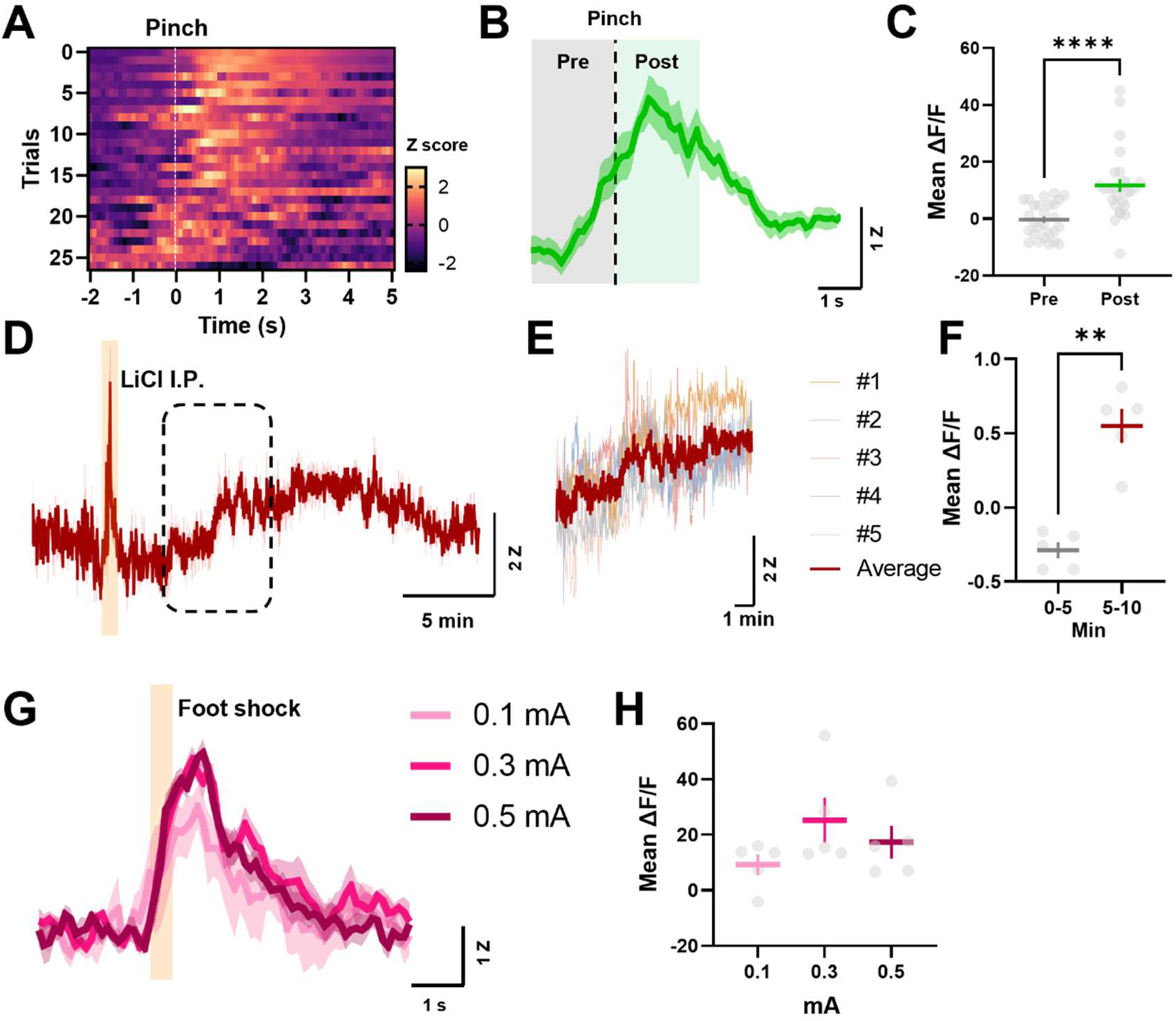
PBN *Ntsr1* neuronal activities in response to aversive stimuli. (A) Heat map of neural activity in response to tail pinch from all mice. White dotted horizontal lines indicate individual mice (*n* = 26 trials in 7 mice). (B) Average trace of neural activities during tail pinch (*n* = 7 mice). (C) Mean fluorescence during the pre 2 s and post 2 s of tail pinch (*n* = 26 trials in 7 mice, paired two-tailed Student’s t-test; *p* < 0.0001). (D) Average trace of neural activities in response to LiCl administration. Yellow box indicates approximate time of IP injection (*n* = 5 mice). (E) Magnified view of individual and average traces corresponding to the dotted box in d). (F) Mean fluorescence during 0-5 min and 5-10 min after LiCl administration (*n* = 5 mice, paired two-tailed Student’s t-test; p = 0.0021). (G) Average neural activities in response to various intensity of electric foot shock. Yellow box indicates shock duration (*n* = 5 mice). (H) Mean fluorescence in response to different shock intensities (*n* = 5 mice, One-way RM ANOVA). Data are presented as mean ± SEM.

**Fig. S8.**
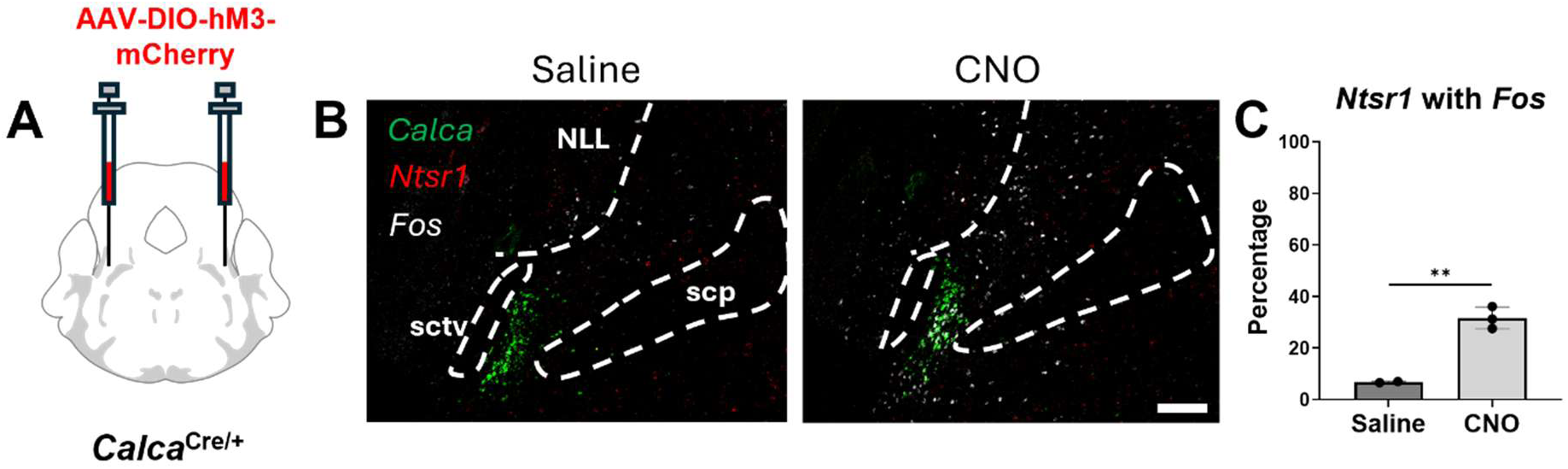
PBN *Calca* neuronal activation results in *Fos* expression in PBN *Ntsr1* neurons. (A) Schematic of viral injection of AAV_DJ_-DIO-hM3Dq-mCherry into the PBN of *Calca^Cre/+^* mice (B) Representative image of expression pattern of *Calca*, *Ntsr1*, and *Fos* in the PBN Saline or CNO administration (C) Activation of PBN *Calca* neurons with CNO increased *Fos* mRNA in PBN *Ntsr1* neurons (*n* = 2 PBN sections for saline, *n* = 3 PBN sections for CNO, unpaired two-tailed Student’s t test; p<0.0045) Scale bar: 200 µm. Data are presented as mean ± SEM.

**Fig S9.**
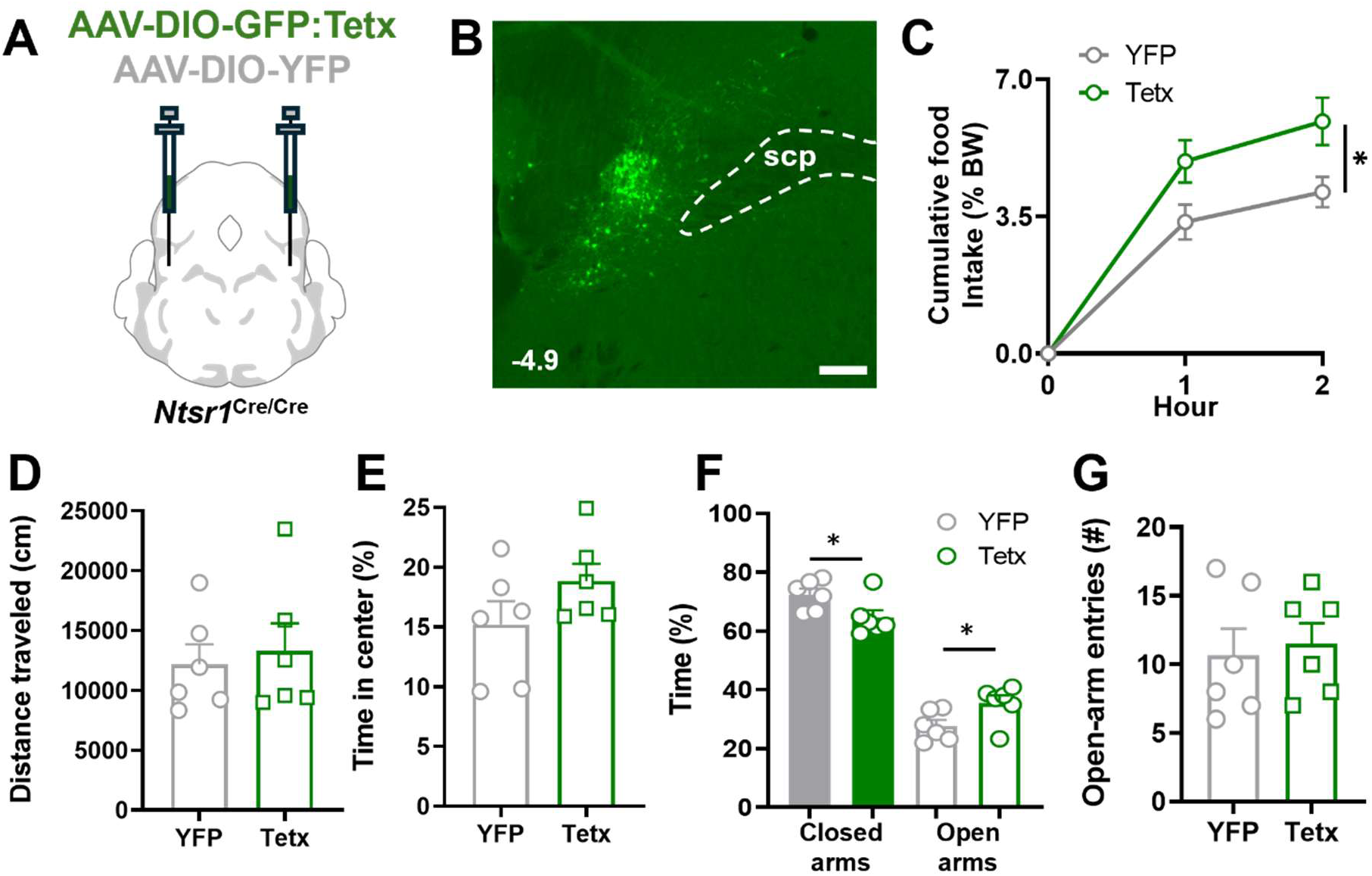
Silencing Ntsr1 neurons increases food intake and reduces anxiety. (A) Schematic of viral injection of AAV-DIO-GFP:Tetx into the PBN of Ntsr1Cre/Cre mice. (B) Representative image of Tetx expression in PBN. (C) Cumulative food intake calculated as percent body weight (Two-way RM ANOVA; *p* = 0.0341 for YFP vs Tetx). (D) Distance traveled in the open field test (Unpaired two-tailed Student’s t-test; *p* = 0.6999). (E) Percent time in center during open field (Unpaired two-tailed Student’s t-test; *p* = 0.1645). (F) Percent time spend in closed and open arms in the elevated plus maze (Student t-test; *p* = 0.0386 for closed arms and p = 0.0385 for open arms). (G) Total number of open arm entries in the elevated plus maze (Unpaired two-tailed Student’s t-test; *p* = 0.7399). Scale bars, 200 µm, n = 6 for YFP and n = 6 for Tetx mice. Data are presented as mean ± SEM.

**Fig. S10.**
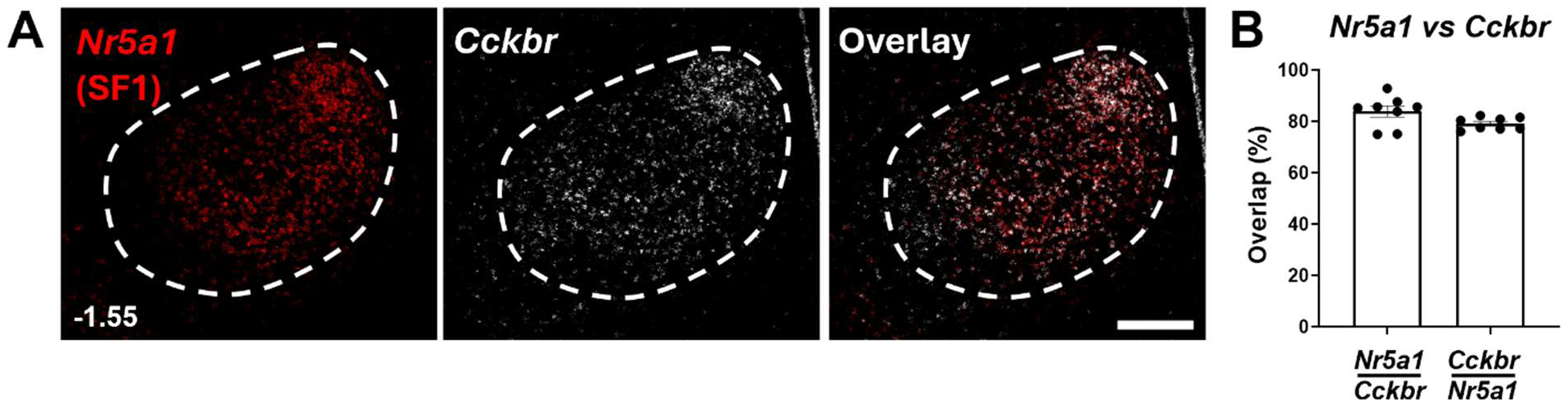
*Nr5a1* neurons in the VMH coexpress *Cckbr*. (A) Representative images of *in situ* hybridization for *Nr5a1* (red), *Cckbr* (white). (B) Percent overlap between *Cckbr* and *Nr5a1* positive populations in the VMH (*n* = 8 whole VMH for both groups) Scale bars: 200 µm. Numbers in lower left indicate bregma level. Data are presented as mean ± SEM.

